# Virus ΨM2 enters a non-lytic lifecycle alternative to lysogeny during infection of *Methanothermobacter marburgensis*

**DOI:** 10.64898/2026.10.07.757236

**Authors:** Sylvia M. Schlaich, Josephine Herbert, Donat Crippa, Beyza R. Kercin, Marie C. Schoelmerich, Bastian Molitor

## Abstract

Methanogens, a diverse group of archaea that produce methane during energy metabolism, are prevalent in anoxic habitats where they are exposed to viral infections. The head-tailed virus ΨM2 is one of few virus isolates that infect a methanogen host and was described as a lytic virus that lyses its host, *Methanothermobacter marburgensis*. Here, we present evidence that ΨM2 does not exclusively follow the classic lytic lifecycle but can also enter an alternative, chronic infection-like state. Growth experiments showed that infected cultures in minimal medium, optimized for the methanogen host (MS medium) did not exhibit complete lysis but continuously released infective virions. In contrast, in a suboptimal complex medium (SAB medium), the ΨM2-infected culture released virions in the exponential growth phase and eventually lysed after 14 hours post infection. Although the genome of ΨM2 encodes a putative integrase, a prerequisite for lysogeny, whole genome sequencing of ΨM2-infected *M. marburgensis* cultures revealed that the viral DNA remained extrachromosomal rather than integrating into the host genome, likely due to a mutation in the integrase gene. These findings suggest that ΨM2 enters a non-lytic lifecycle alternative to lysogeny under suboptimal host growth conditions. Sequencing furthe identified a heterogeneous viral population including previously undescribed variants. A high density of non-synonymous mutations and frameshift-inducing insertions was observed within ORFs predicted to encode a capsid protein and a major spike protein, suggesting these viral variants to differ in host recognition and infectivity. Overall, our results show more complex viral behavior and greater genetic variability than previously described.

**Importance:** Methanogens directly contribute to global greenhouse gas emissions, but they can also be used to mitigate carbon dioxide emissions for biofuel production. Furthermore, they are increasingly recognized as important components of the human microbiota. Understanding the biology of viruses would greatly benefit climate change mitigation strategies as well as research on human gut methanogens. However, only few methanogenic viruses have been isolated to date. One reason might be that lysogeny, which entails viral integration in the genome of their methanogen host, is prevalent but not well understood, and thus, hinders the discovery of novel viruses. In support of this hypothesis, we provide evidence that an archaeal virus previously described as lytic presents an alternative non-lytic lifecycle, which hinders virus discovery through commonly employed lysis-based methods. Research on head-tailed methanogen viruses may provide insights into common features and potential evolutionary connections with the well-studied group of head-tailed bacteriophages.

## Introduction

Methanogenic archaea (methanogens) generate methane as the final product of their metabolism. Hydrogenotrophic methanogens, in particular, convert molecular hydrogen and carbon dioxide into methane. This confers them a unique ecological role as both consumers and producers of different greenhouse gasses. In their role as greenhouse gas producers, they contribute to the rising global methane emissions into the atmosphere, mainly from wetlands and the guts of ruminants for milk and meat production (Filonchyk et al., 2024; Rosentreter et al., 2024). In the opposing role, methanogens are utilized in biotechnological applications to capture the greenhouse gas carbon dioxide for the controlled production of methane as renewable biogas (Contreras et al., 2022; Enzmann et al., 2018; Martin et al., 2013). Additionally, methanogens are recognized as important components of the human gut microbiome, with limited understanding of their role (Chibani et al., 2022; Dridi et al., 2011; Kuehnast et al., 2025). Research on hydrogenotrophic methanogens suffers from general limitations due to their anaerobic, gas-dependent cultivation conditions (Sun et al., 2020), as well as expanding but limited availability of tools for the targeted manipulation and investigation of these microbes (Costa & Whitman, 2023; Klein et al., 2025; Mühling et al., 2024).

One underexplored way to study and manipulate methanogens in their diverse habitats relies on the use of viruses that infect them either as predators to eliminate specific species from a microbial community or as vectors to deliver genetic material to their microbial hosts. The biggest challenge to the approach of manipulating methanogen populations by viruses is the prevailing scarcity of research on viruses that infect these microbes. Most isolated archaeal viruses infect extremophilic species of the genera *Saccharolobus* and *Halorubrum*, which host an impressively diverse virosphere (Krupovic et al., 2018; Prangishvili et al., 2017). Some archaeal viruses seem related to known bacteriophages and show many common features, while other archaeal viruses display peculiarities that cannot be found in bacteriophages and viruses of eukaryotes (Krupovic et al., 2018; Prangishvili et al., 2017). Interesting unique features range from new virion morphologies (Pietilä et al., 2014) to special virion egress mechanisms (Bize et al., 2009).

Nine functional viruses of methanogens have been described to date, eight of which are head-tailed viruses, namely MSTV1, infecting *Methanobrevibacter smithii* (Baquero et al., 2024), Drs3, infecting *Methanobacterium formicium* (Weidenbach et al., 2021), Blf4, infecting *Methanoculleus bourgensis* E02.3 (Weidenbach et al., 2021), MFTV1, infecting *Methanocaldococcus* spp. (Thiroux et al., 2021), spherical virus MetSV, infecting *Methanosarcina* spp. (Weidenbach et al., 2017), ФF1 and ФF3, infecting different *Methanothermobacter* spp. (Nölling et al., 1993), and ΨM1 and ΨM2, infecting *Methanothermobacter marburgensis* (Meile et al., 1989; Pfister et al., 1998). While MSTV1 and MFTV1 were found in the so-called lysogenic lifecycle, that is, integrated in their host’s genome as proviruses, all other mentioned head-tailed methanogen viruses were only described as lytic, meaning they lyse their hosts to release progeny virions and spread. However, metagenomic studies suggest that there is a vast viral community associated with methanogens, with many putative proviruses identified in methanogen genomes (Medvedeva et al., 2023). While the lysogenic lifecycle can be beneficial to viruses as well as their hosts, we hypothesize that it constitutes a main hindrance to methanogen virus isolation, investigation, and utilization. ΨM2, which infects *M. marburgensis*, was obtained from prolonged propagation of the original isolate, ΨM1, which was isolated from an anaerobic sludge digester (Meile et al., 1989). Compared to the ΨM1 genome, the ΨM2 genome carries a 692 bp deletion. A similar virus genome was discovered integrated into the genome of *Methanothermobacter wolfeii*, called ΨM100, and found to be unable to produce active virions, but still led to autolysis of its host upon hydrogen starvation (Luo et al. 2001). Two studies described ΨM2 as lytic and provided genome data, host growth and infection curves, and an electron micrograph of the head-tailed virion (Meile et al., 1989; Pfister et al., 1998). It was seen that while the open reading frame (ORF) 29 was annotated as an integrase, no lysogenic stage was observed, possibly due to a loss of function mutation in the integrase gene (Pfister et al., 1998). In this study, we revisit the ΨM2 lifecycle. We provide evidence that this virus is not purely lytic but under certain conditions persists as an episome without fully lysing its host. Sequencing results further reveal a virus-genome population composed of ΨM1, ΨM2, and previously undescribed variants. In combination with the development of methods to genetically manipulate the *M. marburgensis* host (Klein et al., 2025; Unger et al., 2026), and a closely related species, *Methanothermobacter thermautotrophicus* (Fink et al., 2021; Fink et al., 2023; Baur et al., 2026), we argue that ΨM2 can serve as a model system to study methanogen viruses and potentially viral lifecycle decisions.

## Results

### ΨM2 establishes a persistent infection in liquid MS medium

To analyze infection dynamics between ΨM2 and *M. marburgensis*, a previous study had only used high-titer infection assays with a multiplicity of infection (MOI) of 3.75 (Meile et al., 1989). In these one-step growth experiments, the infected culture stopped growing from the time point of infection. The latent period, which is the time it takes for lysis to occur, was four hours (Meile et al., 1989). Here, we performed one-step growth experiments with a MOI of 0.1 in the standard medium for *Methanothermobacter* spp. cultivation, MS medium (Balch et al., 1979; Fink et al., 2021), which closely resembles the medium that was used by Meile et al. (1989). Under these conditions, we observed no lysis of the ΨM2-infected *M. marburgensis* cultures (**Figure 1 A**). The uninfected control cultures reached a final optical density (OD_600_) of 0.393 ± 0.053, while the ΨM2-infected cultures reached a final OD_600_ of 0.361 ± 0.077. The doubling time in MS medium did not differ significantly between the uninfected control cultures and the infected cultures, with 2.6 ± 0.2 h versus 2.6 ± 0.3 h (*t*(6) = 0.042, *p* = 0.96). Right after infection of the culture, 8.4·10^5^ ± 9.4·10^5^ plaque-forming units (PFU)/mL were measured (**Figure 1 B**). The number of free virus particles then dropped to 3.1·10^3^ ± 1.2·10^3^ PFU/mL after four hours, which was likely due to adsorption to the host cells. Even without apparent lysis, the infected *M. marburgensis* culture showed a steady increase in free virus particles throughout the exponential growth phase. After 24 hours, the amount of measured free virus particles was 1.3·10^7^ ± 1.1·10^7^ PFU/mL, which represents a ∼15-fold increase of the viral concentration that was administered for infection.

**Figure 1.**
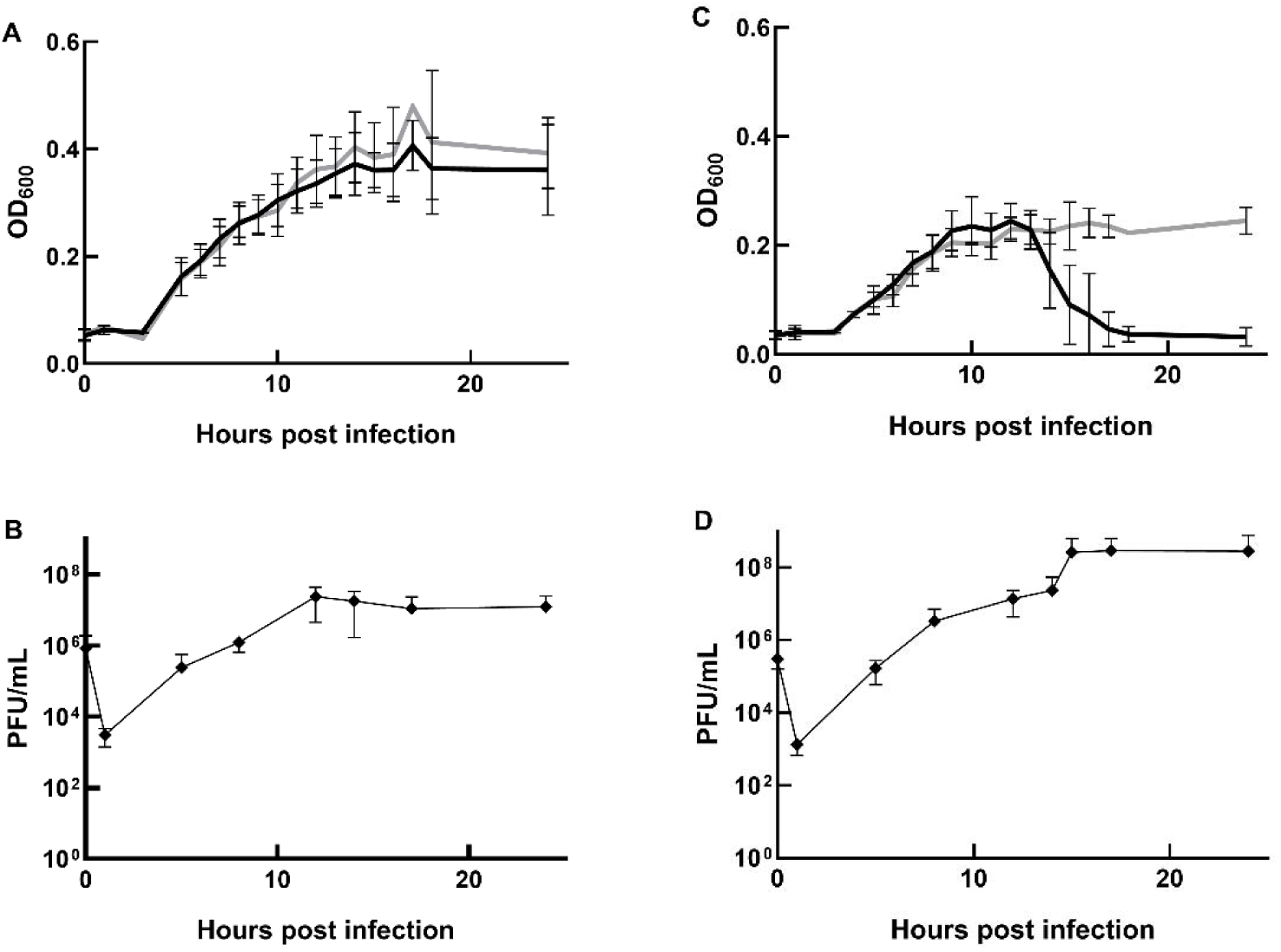
Infection assays comparing infected (black) and uninfected (grey) cultures in MS medium (A) and SAB medium (C). Free virus titer was measured in infected cultures in MS medium (B) and SAB medium (D) and shown as plaque-forming units (PFU)/mL over time (hours post-infection). Error bars indicate the standard deviation of three biological replicates.

### ΨM2 lyses its host at the onset of stationary phase in liquid SAB medium

We observed a different infection outcome in complex growth medium (SAB medium) (Khelaifia et al., 2013). This medium was developed for the cultivation of various methanogen species and generally differs from MS medium by slightly increased concentrations of salts, as well as the presence of additional components, such as yeast extract, acetate, and formate (Materials and Methods). While the uninfected control cultures reached a final OD_600_ of 0.272 ± 0.019, the ΨM2-infected cultures lysed after a latent period of approximately 14 hours post infection after reaching an OD_600_ of 0.245 ± 0.030 (**Figure 1 C**). The doubling time in SAB medium did not differ significantly between the uninfected control cultures and the infected cultures, with 2.4 ± 0.3 h versus 2.2 ± 0.1 h (*t*(7) = 0.819, *p* = 0.44). However, the mean doubling time in all SAB medium cultures was 2.3 ± 0.2 h, and thus, significantly shorter than the mean doubling time of 2.6 ± 0.3 h in all MS medium cultures (*t*(15) = 2.449, *p* = 0.027). The analysis of free virus particles showed the same initial drop in virus numbers as in MS medium, from 3.0·10^5^ ± 1.0·10^5^ PFU/mL directly post infection to 1.3·10^3^ ± 5.7·10^2^ PFU/mL due to adsorption after one hour (**Figure 1 D**). Similar to the observation for MS medium, we found a continuous increase in free virus particles during exponential growth of the ΨM2-infected *M. marburgensis* in SAB medium. Around the timepoint of lysis, the free virus number leaped from 2.4·10^7^ ± 2.8·10^7^ PFU/mL to 2.6·10^8^ ± 3.2·10^8^ PFU/mL, at which level it remained for the rest of the experiment. In SAB medium, the number of free virus particles that we measured throughout the infection assay experienced almost a 1000-fold increase, and the maximum amount of free virus particles was approximately 20-times higher than during infection of *M. marburgensis* in MS medium. Additional experiments on MS medium plates showed that yeast extract, a component of SAB medium, seemed to impact the virus-host relationship resulting in differences in the virus plaque appearance (**Supplementary Results 1**, **Supplementary Figure 1**).

### Whole genome sequencing reveals no integration of the ΨM2 genome into the host genome

During the analysis of MS medium plates from plaque assays, we observed the formation of *M. marburgensis* colonies within plaques 24 h after ΨM2 infection (**Supplementary Result 1, Supplementary Figure 2**). This is the typical appearance of lysogens on plates (sometimes called “mesa”) (Altamirano & Barr, 2021). To test whether the observed differences in infection outcomes and “mesa” were due to the virus entering a lysogenic state, cultures of *M. marburgensis* were infected with ΨM2 and incubated until early exponential (5 hours post inoculation), early stationary (9 hours post inoculation) and late stationary growth phase (24 hours post inoculation) in MS and SAB liquid medium. In both media, we confirmed successful infection of *M. marburgensis* with ΨM2 (**Supplementary Figure 3**). We performed whole genome sequencing from all samples, which we refer to as A-G, corresponding to sample A (MS medium, early exponential timepoint), B (MS medium, early stationary timepoint), C (MS medium, late stationary timepoint), D (SAB medium, early exponential timepoint), E (SAB medium, early stationary timepoint), F (SAB medium, late stationary timepoint), as well as a control sample G (MS medium, early exponential timepoint) from an uninfected *M. marburgensis* culture.

The sequencing run generated 27 Gb of data consisting of 10.31 million reads with an estimated N50 of 3.76 kb (**Supplementary Table 1**). All samples were successfully sequenced, except for sample C, where gDNA extraction had yielded less than 60 ng total gDNA. All other samples were used to assemble genomes in the expected *M. marburgensis* genome size of 1.6 Mbp using *de novo* assembly. The ∼26.9 kb genome of ΨM2 was recovered from reads in all infected samples grown in SAB medium (samples D, E, and F), as well as from the infected MS medium sample at the early exponential timepoint (sample A). As expected, no contigs corresponding to ΨM2 were detected in the uninfected *M. marburgensis* control (sample G), which contained only the host genome. Unexpectedly, initially no reads or contigs matching ΨM2 were recovered from sample B (infected MS medium, late stationary phase). However, subsequent read-mapping analyses identified reads aligning to the ΨM2 genome in this sample, indicating that the virus was present but not recovered during the initial assembly or contig identification step.

Thus, all infected samples yielded three contigs (**Supplementary Table 2**). These three contigs consisted of the contig identified as the *M. marburgensis* genome (1.6 kbp), a circularized contig of 32,360 ± 12,049 bp in length, and finally one smaller circularized contig of 9,514 ± 3,994 bp in length. The second largest circularized contig (32 kbp) was identified in all assemblies except for the control (sample G). This contig matches the expected 26,972 bp length of the YM2 virus genome, with the exception of sample B, where the size was 53,915 bp. Quast analysis further confirmed that this viral contig in sample B did not align with the *M. marburgensis* reference genome (**Supplementary Table 2**). The smaller (9.5 kbp) circularized contig, which was present across all samples, was identified as the pME2001 plasmid (4,439 bp), which is a known component of *M. marburgensis.* BLAST analysis confirmed this identification, showing >99.9% identity across the entire sequence (**Supplementary Table 3**).

Consistent with these findings, geNomad and VIBRANT identified the second largest contig (32 kbp) as viral (excluding sample B) (**Supplementary Tables 4 and 5**). GeNomad classified these contigs within the *Leisingerviridae* family, to which ΨM2 belongs, containing 40-41 total genes, 5-6 hallmark genes, and a virus score of 0.990 ± 0.001 (FDR: 0.004 ± 0.002). VIBRANT yielded similar results, identifying the same viral contigs and gene counts, with v-scores (reflecting the degree of viral association of the predicted annotations) of 7.32 ± 0.00 for KEGG, 17 ± 1.42 for Pfam, and 32.41 ± 4.99 for VOG HMM profiles. Finally, a BLAST search using the ΨM2 genome (Accession No.: GCF 000837485) resulted in alignments to these same contigs (32 kbp, except for sample B) at 99.78 ± 0.55% identity across alignment lengths ranging from 54 to 14,139 bp (3,115 ± 4,243 bp; E-value: 7.09·10^-23^ ± 1.32·10^-22^) (**Supplementary Table 6**). Notably, no viral genes or sequence similarities were detected within the main *M. marburgensis* genome contig. This indicates that although the virus is clearly present within the samples, the ΨM2 genome is not integrated into the host genome.

### Evidence of heterogeneous ΨM1 and ΨM2 viral populations and undescribed variants

Analysis of the high-quality mapped viral reads revealed a 692 bp insertion at position 22,472, which was identified as the DR1 element that was present in the parental ΨM1 virus, and that was considered to have been lost in ΨM2 due to homologous recombination (Pfister *et al*., 2002). The presence of this insertion across all samples at frequencies of 18.96 ± 2.08% suggests a mixed viral population of ΨM1 and ΨM2 types, with ΨM2 being the dominant type (**Supplementary Table 7**).

In addition to this known and characterized insertion, several further genomic variants were identified relative to the published ΨM2 reference sequence (Accession No.: NC_001902.1), most of which have not been previously described (**Figure 2A, Supplementary Table 8**). These include a large insertion/duplication (∼124 bp) in an intergenic region at position 2,622. This was present in all samples at a frequency of 32.64 ± 3.99% and may represent a tandem duplication event. Immediately downstream, a single base pair insertion was detected within a predicted ORF encoding an ASCH domain-containing ribonuclease, which is implicated in RNA metabolism in *E. coli* (Meng *et al*. 2025). This insertion induces a frameshift that changes the amino acid sequence, likely rendering the putative enzyme dysfunctional. At position 16,608 of the genome, within a tandem repeat region previously identified by Pfister *et al*. (2002), we identified insertions of varying lengths at a frequency of 62.32 ± 3.15%.

**Figure 2.**
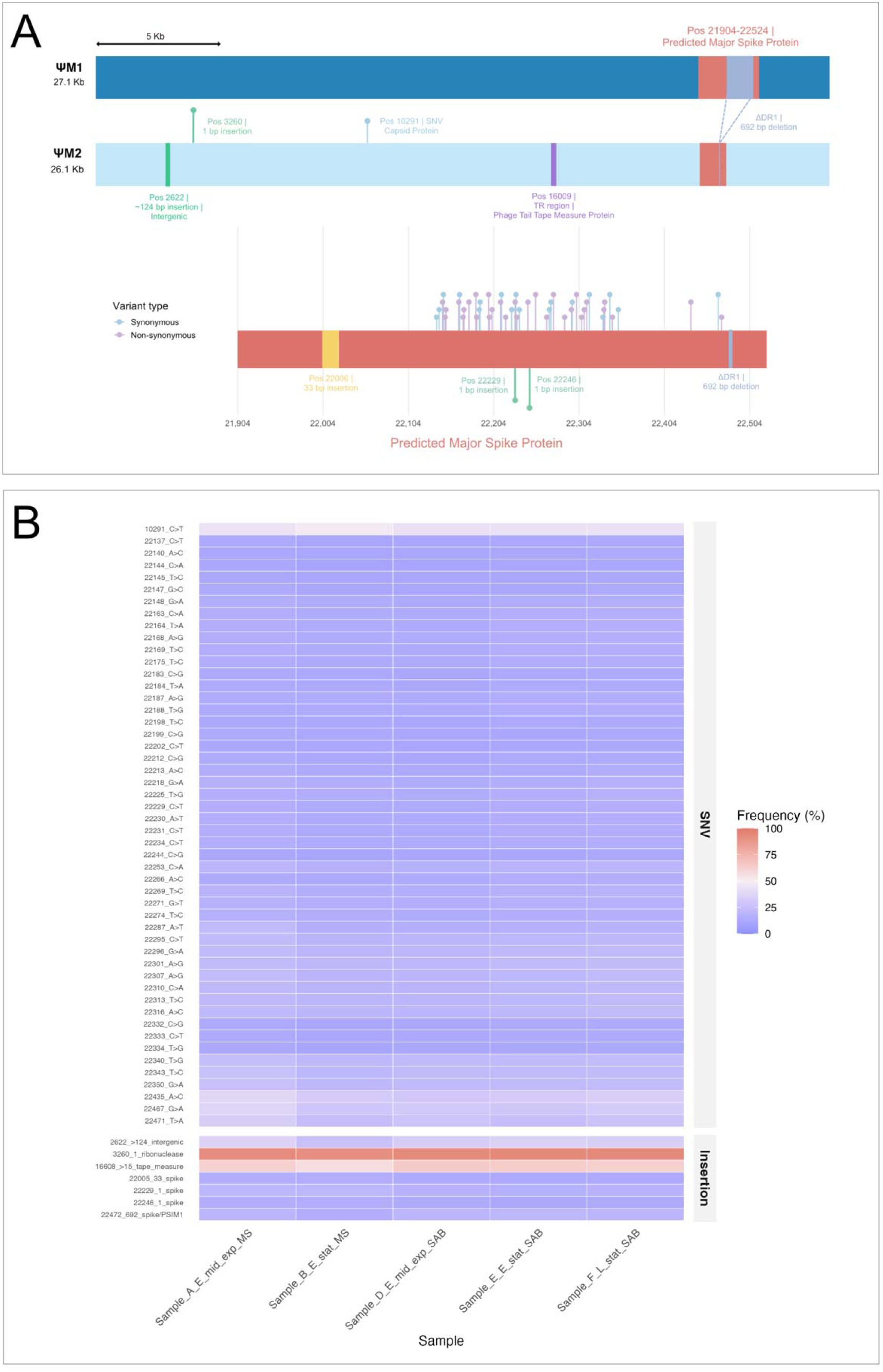
(A) Top: schematic representation of variants identified in this study relative to the NCBI ΨM2 reference genome (Accession No.: GCF 000837485). Detected variants include insertions and single-nucleotide variants (SNVs) across the whole genome. Bottom: zoomed view of the ORF where most variants occur, which is predicted to encode the predicted major spike protein. (B) A heatmap showing the frequency (%) of each identified variant (SNVs and insertions) across all sequenced samples.

Most variants were identified within a previously unannotated open reading frame (ORF) (positions 21,904-22,524), which is now predicted to encode a major spike protein, a crucial structure for viral infection and cell entry. In total, 50 single nucleotide variants (SNVs) were identified at positions common across all samples, with variant allele frequencies of approximately 16.50 ± 4.93%. Notably, 26 of these SNVs resulted in amino acid substitutions. In addition, a 33 bp insertion (frequency of 14.10 ± 1.13%) at position 22,005 was identified, as well as two 1 bp insertions at positions 22,229 (18.26 ± 1.56%) and 22,246 (13.86 ± 1.44%). While these insertions carry frameshift potential that could significantly impact protein structure, their lower frequencies suggest they are not a major subtype.

One SNV was identified at position 10,291, within a predicted capsid protein ORF. This variant occurred at the highest frequency (44.54 ± 2.05%) but was identified as synonymous. The presence of SNVs in only two ORFs, both encoding structural components, may hold significance for viral infectivity and warrants further investigations. Variant frequencies were highly consistent across samples and showed no clear enrichment associated with growth phase or medium (**Figure 2B, Supplementary Table 8**). Consequently, no link between these variants and potential phenotypic effects can be inferred from this dataset.

### Similar virus adsorption is observed in both media

To determine whether differences between the media impact adsorption of the viral particles, we performed more detailed adsorption assays. Within the first hour after infection, we observed a decrease in free virus particles from 2.9·10^4^ ± 8.0·10^3^ PFU/mL to 1.9·10^4^ ± 6.6·10^2^ PFU/mL in MS medium and from 2.8·10^4^ ± 2.6·10^3^ PFU/mL to 1.8·10^4^ ± 4.3·10^2^ PFU/mL in SAB medium (**Figure 3**). Afterwards, the numbers in MS medium fluctuated around a similar level, before rising at the four-hour timepoint (2.5·10^4^ ± 1.5·10^3^ PFU/mL). In SAB medium, we observed a similar fluctuation, but a lower number of free virus particles was reached at 4 hours after infection (1.4·10^4^ ± 2.9·10^3^ PFU/mL). Additionally, adsorption assays with a longer duration showed that the numbers of free virus particles increased strongly after 4 hours post infection with a clearer exponential rise in SAB medium compared to more fluctuating numbers in MS medium (**Supplementary Figure 4**). The adsorption rate constant was 3.8·10^-7^ ± 2.0·10^-7^ mL min^-1^ in MS medium and 3.5·10^-7^ ± 9.7·10^-8^ mL min^-1^ in SAB medium (**Equation 1** in Materials and Methods, **Supplementary Table 9**). Transmission electron microscopy (TEM) analysis of ΨM2-infected *M. marburgensis* cultures confirmed successful adsorption in both media and further revealed that the viral particles attach to the poles of the host cells using their tails (**Figure 4A, B**). Samples with a higher virus-host ratio showed an accumulation of virus particles at the host cell poles (**Supplementary Figure 5**). Those images also indicated a connection to *M. marburgensis* cell appendages (**Supplementary Figure 5**). In few instances, virus particles were imaged at random spots on the *M. marburgensis* cell, with either their whole bodies or only their heads in close proximity to the cell wall (**Supplementary Figure 6**). Meanwhile, analysis of isolated virus concentrates confirmed previous observations on the structure and size of ΨM2 viral particles (**Supplementary Table 10**) (Meile et al., 1989).

**Figure 3.**
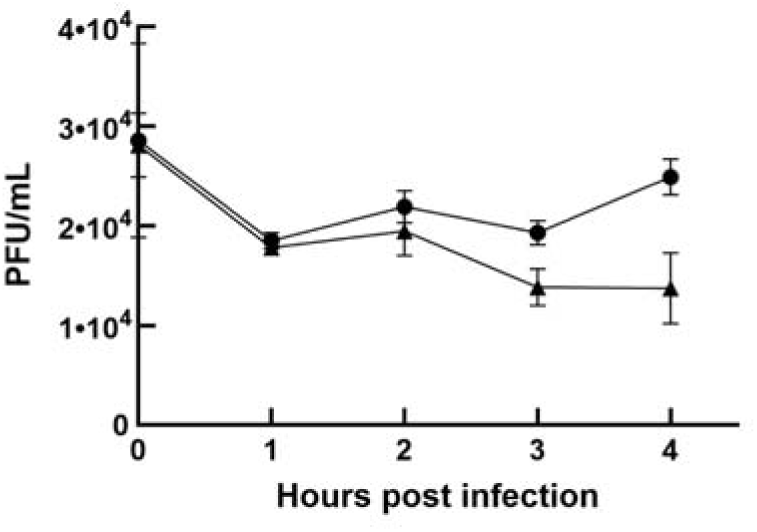
ΨM2 adsorption to *M. marburgensis* in MS medium (circles) and SAB medium (triangles). The concentration of free virus particles (PFU/mL) measured in three biological replicates is plotted against the time in hours post-infection.

**Figure 4.**
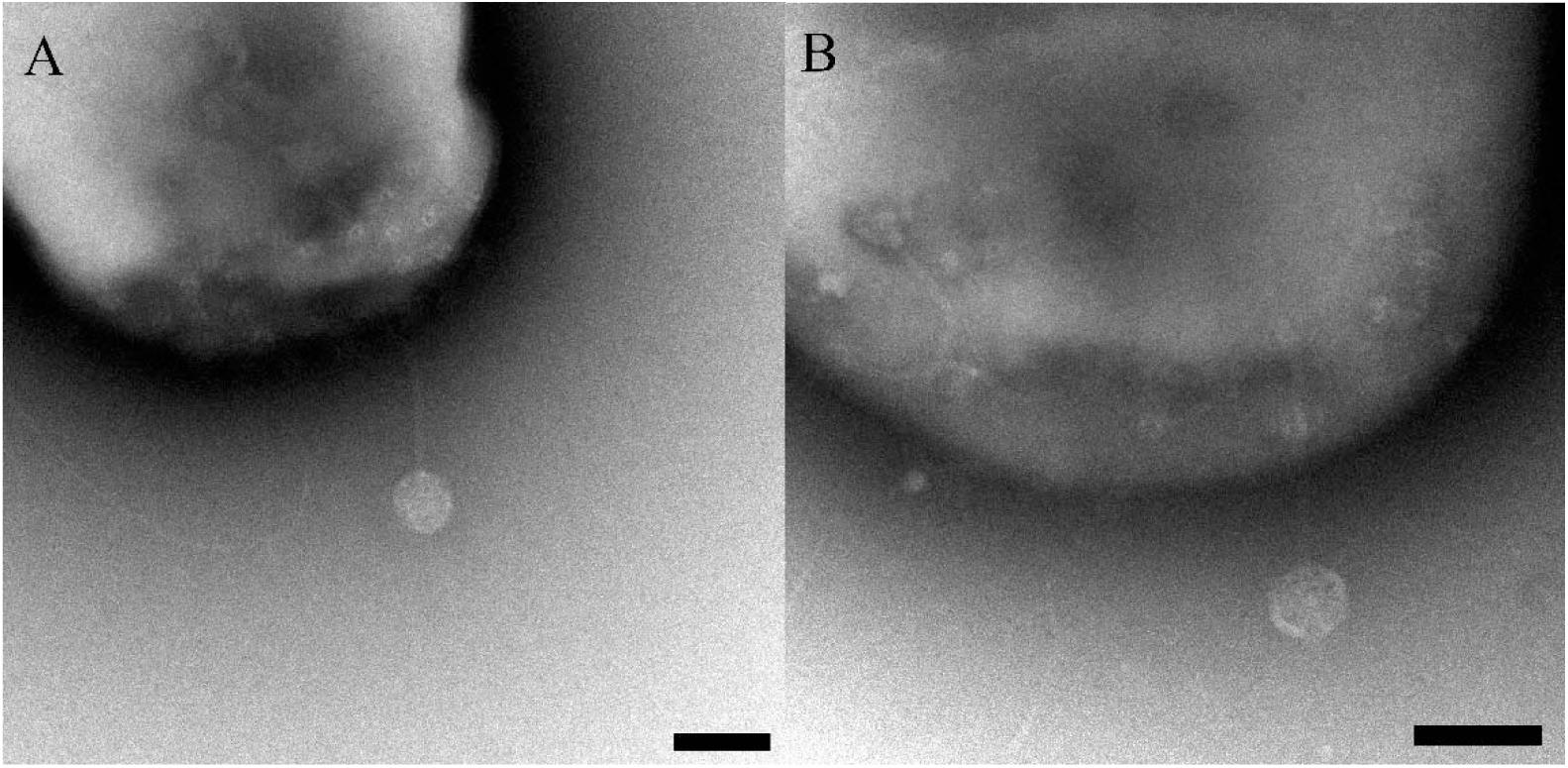
Electron micrographs of ΨM2 virus particles attaching to *M. marburgensis* cell poles in cultures in MS medium (A) and SAB medium (B) at 3.5 h post-infection. The scale bars represent 100 nm.

## Discussion

Here, we provide experimental evidence that the methanogen virus ΨM2, which was considered a lytic virus, undergoes an alternative lifecycle to the lytic one. Our infection assays revealed that not all ΨM2-infected *M. marburgensis* cultures lyse, but that lysis depends on the growth medium used for cultivation. No complete lysis of the culture was observed in minimal MS medium, while complete lysis occurred around 14 hours post infection in complex SAB medium. In both media, however, extracellular virion titers increased from approximately 4 h post-infection. In SAB cultures, this happened before the main lytic event and in MS cultures in the absence of any measurable lysis. Initially, we speculated that ΨM2 is a temperate virus that enters the lysogenic lifecycle when infecting *M. marburgensis* in MS medium. Under this assumption, the observed constant release of viruses throughout the infection cycle, would occur if only subpopulations of the infected cultures were lysed, and the rest remained in the lysogenic lifecycle. This instability of the lysogenic lifecycle has been described before for bacteriophages, including phage lambda, which is one of the best-studied model systems with a lysogenic lifecycle (Little & Michalowski, 2010; Broux et al., 2026). During growth, a low percentage of lysogens (host cells carrying the provirus in their genome) start the lytic cycle without any apparent trigger. Several factors may cause an unstable lysogenic state, including random DNA damage that starts the SOS response, or a sudden decrease in the abundance of lysis-repressor protein during cell division (Little & Michalowski, 2010). Very similar infection dynamics were reported for the lysogenic head-tailed methanogen virus MFTV1 (Thiroux et al., 2021). However, the seldom observation of transitory *M. marburgensis* mesas on plates, which were never observed on plates with incubation times beyond 24 hours, argued against the ability of ΨM2 to enter a stable lysogenic lifecycle. Our sequencing results suggests that ΨM2 does not integrate into the *M. marburgensis* genome, and, thus, the virus does not establish lysogeny; however, we cannot fully rule out that a very small subpopulation carries an integrated virus genome. That no integration occurs is supported by previous findings that the integrase encoded by ΨM2 ORF29 contains a mutation that renders the enzyme inactive (Luo et al., 2001).

Another phenomenon that may explain the observed ΨM2 infection dynamics is pseudolysogeny, in which the viral genome persists within the host in a stalled state, neither replicating productively, nor lysing the cell, until conditions permit restarting the lytic cycle (Mäntynen et al. 2021; Sanchez-Martinez et al., 2025). As the name indicates, this resembles the lysogenic lifecycle, but closer inspection reveals a different mechanism at play. However, this lifecycle is characterized by a lack of released virus particles, whereas ΨM2-infected cultures released virions continuously from approximately 4 h post-infection. Pseudolysogeny, therefore, does not account for our observations as a model for the whole population, but could be considered in a model of a mixed population of many host cells in the pseudolysogenic state while a small subpopulation is being lysed successfully.

Another chronic infection-like phenomenon described in bacteriophages is the so-called carrier-state lifecycle. It resembles lysogeny in that the virus persists within the culture, but there is no lysogen formation through integration of the viral genome. Instead, the infected culture reaches an equilibrium in which a susceptible subpopulation sustains continued progeny virion release through lysis while the rest of the host cells are resistant to the infection and keep up culture growth with little to no growth deficit compared to an uninfected culture (Bastías et al., 2010; Siringan et al., 2014). A carrier-state requires spontaneous switching between cellular resistance and susceptibility. This does not necessarily involve manifested changes in the genome but could be caused by phase variation or growth state-dependent receptor availability.

While our attempts to isolate single colonies from infected but non-lysed cultures never yielded purely ΨM2-resistant cultures, our sequencing results did reveal the presence of a heterogeneous population comprising both ΨM2 and ΨM1, as well as previously undescribed variants (**Supplementary Tables 7 and 8**). Whether these variants were present in the original culture by Pfister *et al*. (1998), and subsequently, the DSMZ stock, or arose during our laboratory passaging, could only be resolved by direct sequencing of the original stock. Variant frequencies remained stable across all samples, giving no indication of selection for or against specific variants. However, as this study was conducted over a single host growth cycle, likely there were insufficient rounds of viral replication for detectable selection to occur. To determine whether one specific variant exhibits increased or decreased infectivity, further experiments involving the passage of the virus over several host growth cycles should be conducted. Notably, the presence of SNVs were confined to two ORFs, the capsid and a predicted major spike protein, both involved in cell adhesion, entry, and virulence (Vinga *et al*., 2012). The presence of 26 non-synonymous variants and 1 bp insertions in each of the two ORFs, could alter protein sequence and structure and may, therefore, influence host interaction, infectivity, and virulence. As both ORFs are relatively small, they may form part of an operon encoding subunits of a larger protein complex. Investigating this would require transcriptomic analyses to identify co-transcribed ORFs, followed by biochemical characterization of the resulting multimeric complexes. Once characterized, targeted viral engineering approaches could be used to assess how the variants observed within this study and resulting structural changes in the encoded proteins influence the infectivity and virulence toward *M. marburgensis*.

From our data, ΨM2 appears to enter a non-lytic, chronic infection state at the onset of infection. In both media, occasional virion release gets triggered, which switches to complete culture lysis only in SAB medium at the beginning of the stationary phase. However, it remains elusive what signal may trigger lysis. The late time point suggests nutrient deficiency, similar to the closely-related defective provirus ΨM100 that is integrated into the *M. wolfeii* genome, which lyses its host upon hydrogen starvation (Luo et al., 2001). In our experiments, hydrogen starvation occurs in both tested media, but only the SAB medium culture lyses upon reaching the stationary phase. This indicates that not hydrogen starvation itself but potentially the general physiological state of the host triggers lysis. Consistent with this, SAB medium is less suitable for *M. marburgensis* growth than MS medium, despite its richer composition, as reflected in lower final OD_600_ in the growth experiments, and we were unable to cultivate *M. marburgensis* on SAB medium plates at all. The growth of methanogens on plates generally appears more difficult than cultivation in liquid medium, which could relate to reduced gas accessibility on solid medium (Fink et al., 2021). This, in turn, is a potential reason, why plaque formation was possible only on MS medium plates, while MS liquid cultures never fully lysed. On MS plates supplemented with yeast extract cell lawns were denser and plaques smaller and surrounded by so-called plaque-halos. In bacteriophages, such halos stem from the depolymerase activity of bacteriophages, which destroys host extracellular polymeric structures involved in biofilm formation (Cornelissen et al., 2012). Biofilm formation has been observed in *Methanothermobacter* spp., especially in bioreactor setups, but remain less studied (Fink et al., 2023; Rademacher et al., 2012). Thus, a potential impact of ΨM2 on biofilm formation of *M. marburgensis* awaits further investigation. Overall, the differences between the media and the two cultivation methods indicate a tendency for ΨM2 to assume a non-lytic lifecycle under optimal host growth conditions and lyse the host cells upon suboptimal host growth conditions.

In more detail, SAB medium differs from MS medium in the presence of yeast extract, trypticase, formate, acetate, and methanol. We assume the factor that facilitates lysis to be either the depletion of a nutrient, trace element, an inhibitory compound, or changes in the metabolism of *M. marburgensis* in response to one (or some) of these additions. For example, *M. marburgensis* cannot utilize formate as an alternative substrate for methanogenesis, but still relies on formate biosynthesis for biomass generation, as this is an important metabolite in purine biosynthesis. The conversion of carbon dioxide to formate is connected to the oxidation of reduced cofactor F_420_ by a formate dehydrogenase (Casini et al., 2023). The presence of formate in SAB medium could impact either the expression of the formate dehydrogenase or the purine and biomass biosynthesis of *M. marburgensis*, which would alter the overall energy metabolism. Formate is, thus, one possible cause of the observed differences in infection outcome between the two media. However, a detailed investigation of single compounds in SAB medium will be required to deduce if a nutrient or a potentially inhibiting substance (*e.g.*, methanol) is involved in the lysis decision of ΨM2.

Another potential lysis trigger could be the virus-host ratio. The original publication (Meile et al., 1989) used a high MOI of 3.75, while we tested a low MOI of 0.1. We observed no signs of lysis in MS medium and no lysis until the stationary phase in SAB medium, while detecting similar concentrations of free virus particles at the beginning of stationary phase in MS medium (1.8·10^7^ ± 1.4·10^7^ PFU/mL, 14 h post infection) and SAB medium (1.4·10^7^ ± 8.2·10^6^ PFU/mL, 12 h post infection). However, the culture density in MS medium (0.373 ± 0.054, 14 h post infection) at this timepoint of the growth experiment was significantly higher than in SAB medium (0.245 ± 0.030, 12 h post infection) (*t*(10) = 4.670, *p* = 0.0009). Thus, the higher apparent MOI could be a factor in the lysis decision of ΨM2. Host cell density in general has been associated with infection outcome for bacteriophages. It has been observed that the lysogenic pathway is especially favorable at low host cell densities, because progeny virions may not find another host cell to infect or eradicate the host population completely, leaving the virus stranded. However, there are also observations indicating that lysogeny is preferred at very high host cell density (Weitz et al., 2017). While the mechanisms are not well understood, there are indications for the importance of density-dependent dynamics in infection outcomes (Cheong et al., 2022).

Additional insights into the infection dynamics of ΨM2 and *M. marburgensis* were generated in adsorption assays. The adsorption rate constant for ΨM2 in either medium was two orders of magnitude higher (3.8·10^-7^ ± 2.0·10^-7^ mL min^-1^ for MS medium and 3.5·10^-7^ ± 9.7·10^-8^ mL min^-1^ for SAB medium) than that of the haloarchaeal virus HFTV1, a head-tailed archaeal virus with fast adsorption (1.9·10^-9^ mL min^-1^) (Schwarzer et al., 2023), and one order of magnitude higher than the adsorption rate of bacteriophage lambda (2.9·10^-8^ mL min^-1^, calculated with Eq. 1 based on data from Shao and Wang (2008)). This constant describes the probability of a single virus to attach to a single microbial cell in a volume of one milliliter per minute. The formula does not consider the temperature, which may lead to an overestimation of how fast ΨM2 adsorption (at 60°C) really is, compared to the other mentioned viruses that infect microbes growing at 37°C. However, the comparatively high adsorption rate constant of ΨM2 can still be interpreted as a high probability of adsorption of the single particles. The timeframe from infection to rising numbers of free ΨM2 particles was the same as in Meile *et al*. (1989). However, the latent period (which was only observed in SAB medium) was almost four times longer in our experiment, likely due to the lower MOI.

ΨM2 particles that were absorbed to the host cell were frequently attached with their tail first, as described for many head-tailed viruses (Schwarzer et al., 2023). However, similar to observations for HFTV1, we observed virus particles in very close proximity to host cells, seemingly binding with other parts of the virion than just the end of the tail. The adsorption of head-tailed viruses is typically a two-step process: initial, reversible binding of the whole virus particle to any site on the host cell or cell appendages, which is followed by irreversible binding of the virus to the host-cell receptor. Thus, our observations reinforce the idea that the step of initial, reversible binding also applies to archaeal head-tailed viruses.

The use of host cell appendages as sites of initial binding has been described in flagellotrophic bacteriophages. The bond in this process will either be made between host pili and the viral head and tail or just the viral head, which prevents the need for detachment of the tail before secondary binding can occur between virus tail and host-cell receptor (Gambino & Sørensen, 2024). The host-cell receptor of ΨM2 is currently unknown. However, it may be a protein located at the cell poles. While *M. marburgensis* is not flagellated, the receptor could be a receptor-type that has also been described for flagellotrophic bacteriophages, such as a pilus portal protein (the membrane-spanning part of the pilus apparatus) or other cell-wall proteins like efflux pumps. Examples of host-cell receptors of bacteriophages that infect Gram-positive bacteria are peptidoglycan and teichoic acids or parts of these macromolecules (Bertozzi 2016). We propose a two-step adsorption model for ΨM2, as described in bacteriophages before. The initial reversible binding in this process may either follow the example of flagellotrophic bacteriophages and happen at a cell appendage (“flagellotrophic adsorption”) or be made between the virus particle and any part of the host cell wall (“tumbling adsorption”) (**Figure 5**). The host-cell receptor is likely located at the cell pole. This model, however, needs to be experimentally investigated. Further tests in liquid medium observing growth behavior and transcriptome differences of the host may bring new insights.

**Figure 5.**
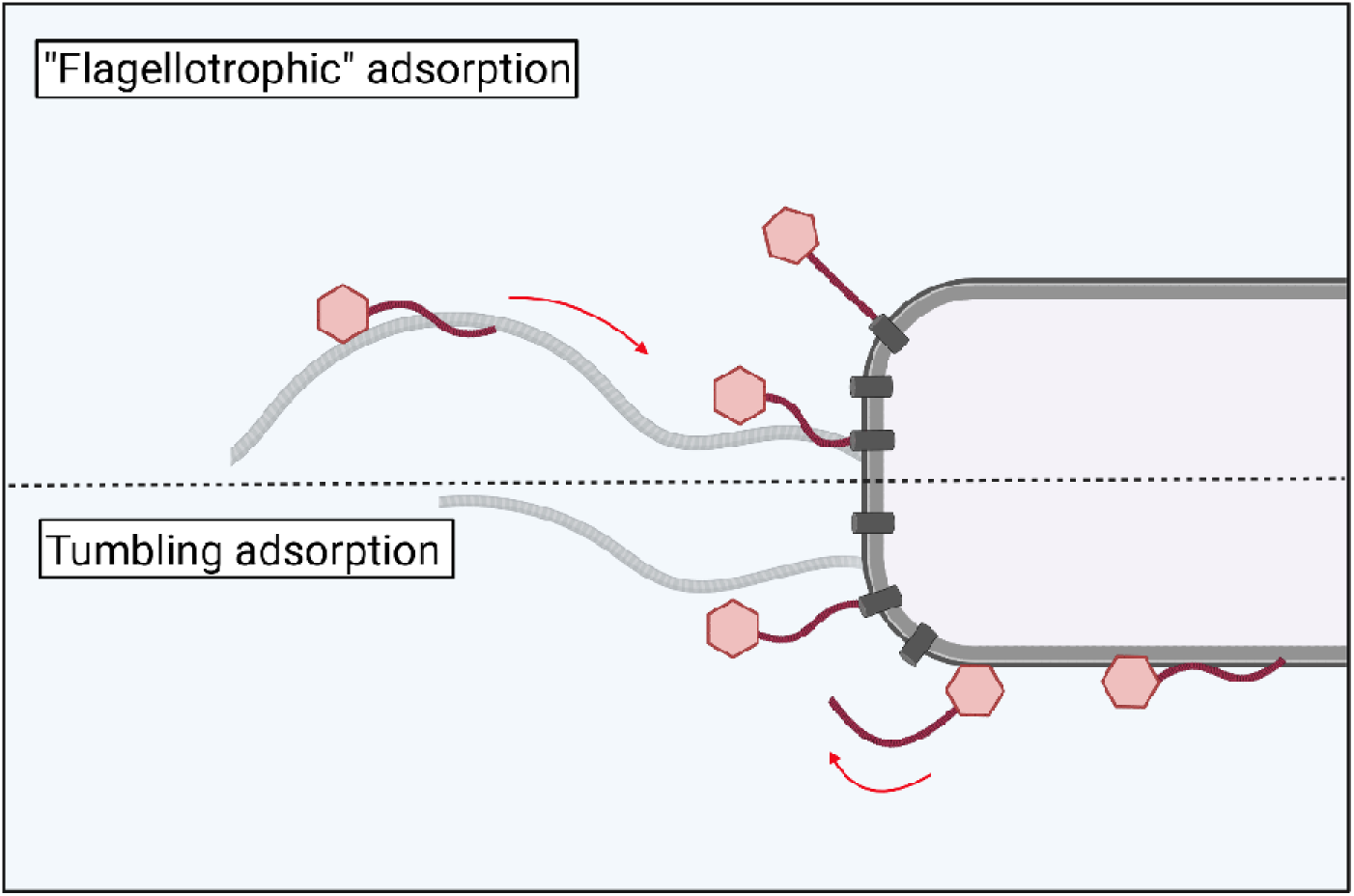
Graphical depiction of the proposed working model for ΨM2 virus adsorption. During “flagellotrophic” adsorption, virus particles (red) reversibly attach to cell appendages, using their heads and tails, and then get transported toward the cell, where they irreversibly bind to their= host-cell receptor (dark grey). During tumbling adsorption, virus particles reversibly bind to th host cell wall, using their heads and tails, and tumble alongside their host through reversible binding and detachment until they find their host cell receptor and bind irreversibly.

In conclusion, our study presents evidence that ΨM2 is not a purely lytic virus but also enters an alternative, chronic infection-like lifecycle under specific host growth conditions. The mutated integrase gene is a strong indicator that the virus does not integrate into the host genome. Potential triggers for lysis are nutrient deficiencies of the host, the virus-host ratio in the infected culture, or a combination thereof. The existence of a heterogeneous ΨM1/ΨM2 population, together with previously undescribed variants, may result in structural and consequently functional changes in key proteins involved in infectivity and virulence, although further studies are required to determine the extent and biological significance of these effects. Based on TEM images, we further hypothesize that ΨM2 may have a potential host-cell receptor located at the cell poles. The identification of the exact host-cell receptor, as well as further investigations on the alternative infection mechanism, will provide a deeper understanding of ΨM2 as a model system for head-tailed methanogen viruses.

## Materials and methods

### Strains, media, and buffers

*M. marburgensis* (DSM 2133) was obtained from the DSMZ (Braunschweig, Germany) and cultivated anaerobically at 60 °C in sealed serum bottles with a headspace of H_2_/CO_2_ (80/20 vol%). The bottles were filled to 20 vol% with the respective growth medium. Anaerobic work was performed in an anerobic chamber (Coy Laboratory Products, Ann Arbor, USA) with a headspace of N_2_/H_2_ (95/5 vol%). For the routine cultivation of uninfected *M. marburgensis,* as well as infection and adsorption assays, MS medium by Balch et al. (1979) was used. This medium contained (per liter): 0.45 g NaCl, 6 g of NaHCO_3_, 0.17 g K_2_HPO_4_, 0.19 g (NH_4_)_2_SO_4_, 0.08 g MgCl_2_·6 H_2_O, 0.06 g CaCl_2_·2 H_2_O, 0.23 g KH_2_PO_4_, 1 mL 0.2 weight% (NH_4_)_2_Ni(SO_4_)_2_, 1 mL 0.2 weight% FeCl_2_·4 H_2_O, 4 mL 0.025 weight% resazurin, and 1 mL 10X trace elements solution. The 10X trace elements solution for MS medium contained (per liter): 30 g MgSO_4_·7 H_2_O, 5 g MnSO_4_·H_2_O, 10 g NaCl, 1 g FeSO_4_·7 H_2_O, 1.8 g CoCl_2_·6 H_2_O, 1 g CaCl_2_·2 H_2_O, 1.8 g ZnSO_4_·7 H_2_O, 0.1 g CuSO_4_·5 H_2_O, 0.18 g KAI(SO_4_)_2_·12 H_2_O, 0.1 g H_3_BO_3_, 0.1 g Na_2_MoO_4_·2 H_2_O, 2.8 g (NH_4_)_2_Ni(SO_4_)_2_·6 H_2_O, 0.1 g Na_2_WO_4_·2 H_2_O, 0.1 g Na_2_SeO_4_. This medium mixture was sparged with N_2_/CO_2_ (80/20 vol%) and brought into an anaerobic chamber, where 0.5 g/L L-cysteine hydrochloride monohydrate was added and the medium was aliquoted into serum bottles. For solid MS medium, 0.3 g/L Na_2_S was also added to the medium before it was aliquoted to serum bottles containing Bacto agar (BD Life Science, Berkshire, United Kingdom) to 1.5 weight% for regular agar or 0.5 weight% for softagar. MS buffer was prepared in the same way as MS medium, but without the trace elements, resazurin, and cysteine. It was further aliquoted in volumes of 5 mL in Hungate tubes (Ochs, Bovenden, Germany) without exchanging the headspace, which was, thus, the same as in the anaerobic chamber. For the propagation of ΨM2 and infection assays, SAB medium by Khelaifia et al. (2013) was used, containing (per liter): 1.5 mg NiCl_2_·6 H_2_O, 0.5 mg FeSO_4_·H_2_O, 0.8 g MgSO_4_·7 H_2_O, 0.5 g KH_2_PO_4_, 0.5 g K_2_HPO_4_, 2 g NaHCO_3_, 0.05 g KCl, 0.05 g CaCl_2_·7 H_2_O, 1.5 g NaCl, 1 g NH_4_Cl, 0.6 mg MnSO_4_·7 H_2_O, 0.1 mg ZnSO_4_·7 H_2_O, 0.02 mg CuSO_4_·5 H_2_O, 0.2 mg KAl(SO_4_)_2_·12 H_2_O, 7 mg H_3_BO_3_, 4 mg CoSO_4_·7 H_2_O, 0.5 mg Na_2_MoO_4_·2 H_2_O, 3 mg Na_2_SeO_3_·5 H_2_O, 4 mg Na_2_WO_4_·2 H_2_O, 0.4 g Na_2_S, 3.24 mL methanol, 0.15 mg nitrilotriacetic acid, 10.8 g sodium formate, 1 g sodium acetate, 2 g trypticase, 2 g yeast extract, 0.5 g L-cysteine hydrochloride monohydrate, 5 mM valeric acid, 5 mM isovaleric acid, 5 mM 2-methylbutyric acid, 6 mM isobutyric acid, 5 mM 2-methyl valeric acid, 1 mg resazurin, and 1 mL vitamin solution (Balch et al., 1979).

### Virus isolation and concentration

For the isolation and concentration of ΨM2 (DSM 12792), which we obtained from the DSMZ (Braunschweig, Germany), cultures of *M. marburgensis* in SAB medium were infected with 200 μL of a virus stock solution and incubated overnight in a 60°C shaking incubator. The lysed culture was then centrifuged at 8,000 rcf and 4°C for 20 min. The cell pellet was discarded and the supernatant sterile-filtered with 0.45 μm syringe filters (Carl Roth, Karlsruhe, Germany). This culture lysate was either stored at 4°C or concentrated further using PEG precipitation. For this concentration method, the culture lysate was mixed with 30 g/L NaCl_2_ and 75 g/L PEG8000 and stirred overnight at 4°C. The next day, the solution was centrifuged at 8,000 rcf and 4°C for 25 min and the supernatant discarded. The pellet was then resuspended in 1% or 10% of the volume of the culture lysate in MS buffer and stored at 4°C. Titer determination of all culture lysates and lysate concentrates was performed with plaque assays.

### Plaque assay

To quantify infective ΨM2 particles from a given sample, plaque assays were performed anaerobically using plates of MS medium. Bottom plates were poured using 9 mL MS agar (1.5%) and dried for about 20 min. A culture of *M. marburgensis* was incubated to an OD_600_ between 0.1 and 0.2 and distributed in 5 mL reaction tubes for each sample to be plated. The virus sample or a dilution thereof was added to one of these tubes, followed by addition of 3 mL of liquefied MS softagar (0.5%). The mix was inverted several times and then pour-plated on one of the bottom plates. For better gas-solid mass transfer of the plates, two sterilized paper clips were attached to the rim of the petri dish. Once dry, the plates were placed into an anaerobic jar upside down and the sealed jar was transferred outside the anaerobic chamber for a gas exchange to 1.5 bar overpressure with H_2_/CO_2_/H_2_S (79.9/20/0.1 vol%). Incubation of the plaque assay was performed in a non-shaking incubator at 60 °C for 3 to 4 days, with a single re-filling of the gas to 1.5 bar overpressure once the pressure dropped to 0 bar.

### Infection and adsorption assays

Infection dynamics between virus ΨM2 and host *M. marburgensis* were studies using two types of growth experiments. For more detailed investigations of the virus lifecycle, infection assays in 50 mL medium bottles were performed. For this, the media were inoculated with *M. marburgensis* to an OD of 0.04 and infected with ΨM2 to a MOI of 0.1. A generated standard curve was used to calculate the number of CFU from OD_600_ of any *M. marburgensis* culture (**Supplementary Figure 7**). The determination of MOI in infection and adsorption assays was then based on the calculated CFU value of the used culture. After 4 h, which is the determined adsorption time of the virus, the cells were washed by centrifugation at 8,000 rcf for 15 min at room temperature and resuspension of the pellet in 5 mL pre-warmed medium. The resuspended pellet was immediately transferred back to a fresh, pre-warmed bottle of medium and incubation continued. Throughout the experiment, including directly after inoculation, 1 mL samples were taken, of which 500 μL were used for OD_600_ measurement and 500 μL were centrifuged at 10,000 rcf, 4 °C for 10 min to collect the supernatant as a viral sample for this timepoint. Viral samples were then analyzed in plaque assays. The infected cultures were prepared in duplicates and uninfected cultures without virus infection were used as controls.

For determination of the adsorption time, *M. marburgensis* was inoculated in 20 mL MS or SAB medium and infected with ΨM2 to a MOI of 0.01. Cultures were then incubated at 60°C and samples for determination of free virus particles *via* plaque assay were taken at different timepoints. As control, medium bottles with the same amount of virus but without host cells were used.

The adsorption rate constant for ΨM2 was calculated from the infected cultures using the equation described in Schwarzer et al. (2023) (**Equation 1**).

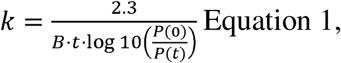

where *k* is the adsorption rate constant, *B* is the host cell concentration, *t* is the time in hours, *P(t)* is the concentration of free virus particles at timepoint *t*.

### Whole genome sequencing

*M. marburgensis* cultures in 50 mL MS and SAB medium were infected with ΨM2 to a MOI of 1. At three different timepoints, namely, at 5, 9, and 24 hours post-infection, triplicate cultures of each medium were sampled. 1 mL was withdrawn for OD_600_ measurement, 1 mL was used for detecting free viral particles (centrifuged at 10,000 rcf and 4°C to store the supernatant at 4°C), and 40 mL of culture were centrifuged at 8,000 rcf for 15 min and the pellet was used for gDNA extraction using a gDNA extraction kit (Macherey-Nagel, Düren, Germany). As a control, three uninfected 50 mL *M. marburgensis* cultures in MS medium were sampled at the early exponential time point of 5 h. The triplicate gDNA samples for each medium and timepoint were pooled and concentrated using a SpeedVac (Thermo Fisher Scientific, Waltham, MA, USA) for 1-2 h at 43°C. After further concentration and clean-up using 2X AMPure XP Beads according to the manufacturer’s protocol (Beckman Coulter, Brea, CA, USA), the final DNA concentrations were measured with the 1X HS dsDNA Qubit assay kit (Thermo Fisher Scientific, Waltham, MA, USA) and the DNA fragment length and integrity with the Tapestation 4150 using the genomic DNA assay (Agilent Technologies, Santa Clara, CA, USA). Sequencing was performed using a PromethION P2 solo (PRO-SEQ002; Oxford Nanopore Technologies (ONT), UK) and the rapid barcoding kit (SQK-RBK114.96) for library preparation following the standard protocol for gDNA recommended by ONT, a negative sequencing control of nuclease-free water was also included within the library. The resulting library was loaded to a R10.1 PromethION flow cell (FLO-PRO114M) and was run for 17.5 h. The raw data was base called using the Super accurate basecalling 400 bps model by Dorado v7.9.8 (ONT) through the MinKnow v25.05.12 (ONT) interface, with parameters including barcode trimming and min. Q-score >8.

### Bioinformatic analyses

The base called raw sequencing data was first concatenated per barcode, followed by Porechop v0.2.4 using default parameters and Chopper v0.10.0b using parameters –minlength 1000 –q 15 for quality control, trimming, and filtering (removing reads with Q<15 and length < 1000 bp). Further stringent filtering was then performed using Filtlong v0.3.1 using parameters --target-bases 500000000 --keep percent 90 --min-length 1000. Quality of the raw and subsequent trimmed and filtered reads was assessed using Nanostat v1.4.0. Flye v2.9 using parameters -- nano-hq --iterations 3 -g 1.75m was used to perform genome assembly using the trimmed and filtered fastq data. The generated genomes were compared with Quast v5.3.0 using the NCBI reference for *M. marburgensis* Marburg as the reference (Accession No.: GCF 027555065.1). GeNomad v1.8.1 was used to identify contigs as host or virus genome, the end-to-end command was used with parameters --cleanup --enable-score-calibration. Integrated virus sequences in the host genome were searched for using VIBRANT v1.2.1 using default parameters and the KEGG KoFam (March 2019 release), Pfam v32 and VOG (release 94) databases. BLAST v2.14.1 was also used to align the NCBI ΨM2 reference genome (Accession No.: GCF 000837485) against each assembly within this study using the parameter --evalue 1e-10 to include only high-quality alignments. Additional analysis included using BLAST to align the NCBI pME2001 sequence (Accession No.: X17205.1) to the assemblies generated within this study using the same parameters as above.

For variant analyses, raw fastq files were mapped to the NCBI ΨM2 reference genome (Accession No.: GCF 000837485) using minimap2 v2.22 using the parameters -ax map-ont -- secondary=no to keep only primary alignments. Alignments were converted to BAM format and filtered using SAMtools v1.16.1 using the command samtools view -F 0x900 -q 20 to remove secondary and supplementary alignments and retain reads with mapping quality ≥20 prior to sorting and indexing. For SNV characterization, the tool InStrain v1.10.0 was used using parameters --min-mapq 10 --pairing_filter non-discordant. For insertion characterization, manual inspection was done using Geneious Prime 2025.2.2, frequencies of the observed insertions were calculated using samtools mpileup with parameters -d 40’000 to account for high coverage and - Q 0 to avoid a second quality filtering step.

Protein function predictions were performed using SeqHub (https://seqhub.org) (Jha *et al*., 2025).

### Transmission electron microscopy

For the visualization of virus particles, transmission electron microscopy was used. Virus and infected culture samples were handed to the Microscopy Facility at the Centre for Molecular Biology of Plants at the University of Tuebingen. There, the samples were incubated for one minute on glow-discharged electron microscopy grids with carbon-coated pioloform film. The grids were then washed on water five times for 10 seconds each. To negatively stain the samples, the grids were then incubated with 1% uranyl acetate for 30 seconds to one minute. Then, the grids were blotted and air dried. Finally, images were captured using a TEM JEM 1400 Plus (Jeol, Tokyo, Japan).

## Data Availability

Genomic sequencing reads reported in this paper are available under SRA Project: PRJNA1529606.

## Acknowledgements

This work was funded by the CMFI Cluster of Excellence in the framework of the DFG (German Research Foundation) under Germany’s Excellence Strategy - EXC 2124 and the Swiss National Science Foundation Starting Grant 226487 (awarded to MSC). We thank Prof. Largus T. Angenent and the Humboldt Foundation in the frame of the Humboldt professorship, which was awarded to LTA, for their support. We further thank the Microscopy Facility at the Center for Plant Molecular Biology at the University of Tübingen for their help with transmission electron microscopy, namely Dr. Sandra Richter, Rebecca Stahl, and Dr. Lorenz Henneberg. We thank Dr. Libera Lo Presti for writing support.

## Author contributions

Sylvia Schlaich (S.S.) and Bastian Molitor (B.M.) conceptualized the study and designed the experiments; together with Josephine Herbert (J.H.), Donat Crippa (D.C.), and Marie C. Schoelmerich (M.C.S.) for the sequencing experiments. S.S. and Beyza R. Kercin (B.R.K.) conducted the growth experiments and infection assays, performed sample preparation for transmission electron microscopy, and together with J.H. and D.C. for genome sequencing. Data analysis was performed by S.S. for growth data, and J.H., and D.C. for sequencing data, under the supervision of B.M. and M.C.S.. S.S. and J.H. prepared figures and tables. S.S. drafted the manuscript, and all authors contributed to editing.

