## Supplementary Material for "Virus ΨM2 enters a non-lytic lifecycle alternative to lysogeny during infection of *Methanothermobacter marburgensis*"

**Supplementary data**

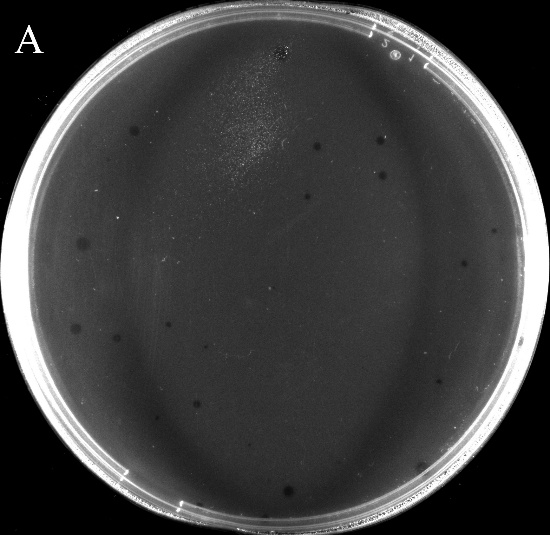

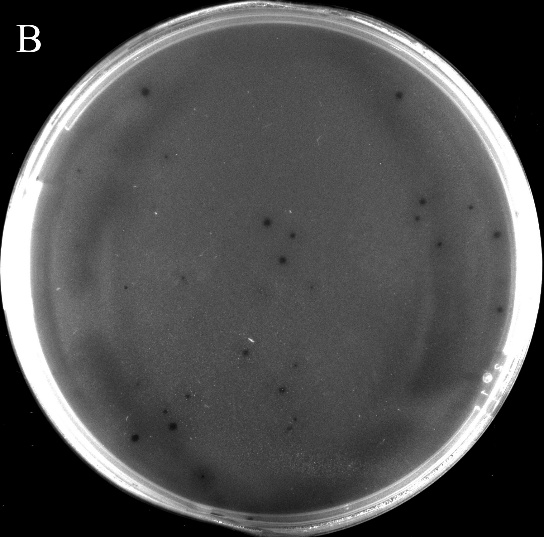

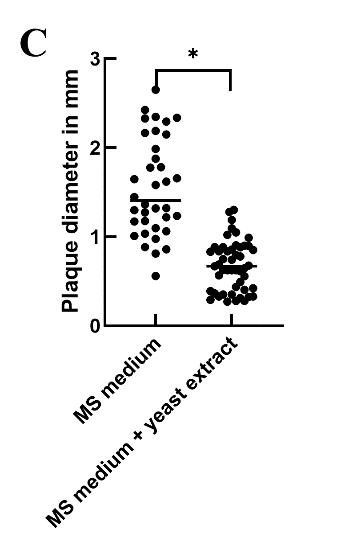

**Supplementary Figure 1.** Plaque formation of ΨM2 on a lawn of M. marburgensis on MS medium (A) and MS medium supplemented with yeast extract (B). The plate pictures were taken in a GelDoc (Analytik Jena, Jena, Germany) and edited for better visibility by increasing brightness and contrast. A graph shows the plaque size analyzed from duplicate plaque assays with and without yeast extract (C). The asterisk indicates a significant difference (p < 0.0001).

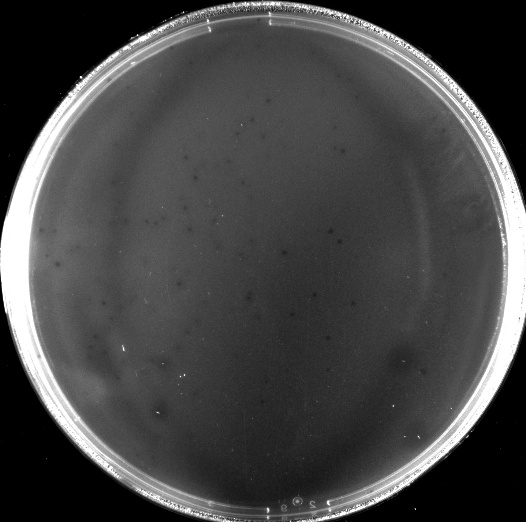

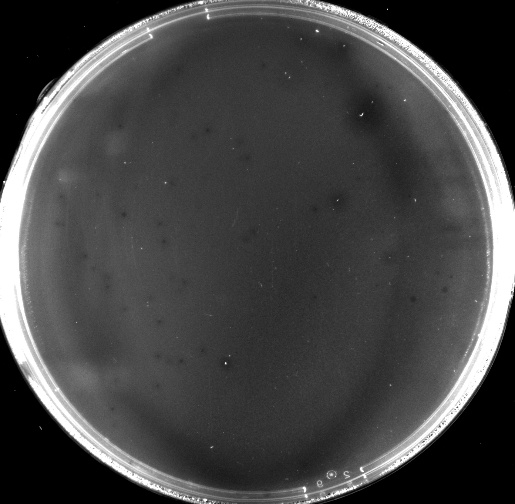

**Supplementary Figure 2.** Pictures of MS medium plates from a plaque assay after ~24h of incubation. The arrows point to wide plaques with host cell colonies in the middle. Images were taken in a GelDoc and edited for better visibility by increasing brightness and contrast.

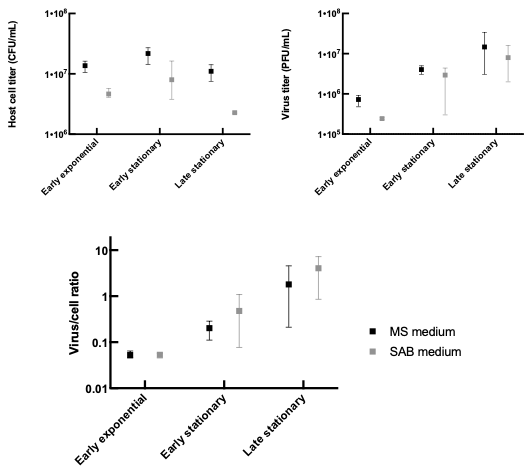

**Supplementary Figure 3.** Growth parameters of ΨM2-infected *M. marburgensis* cultures used for sequencing at three sampling time points (early exponential: 5 hours post-infection (hpi); early stationary: 9 hpi; late stationary: 24 hpi) from triplicate cultures in MS medium (black) and SAB medium (grey). The host cell titer was measured *via* optical density and converted to CFU/mL using a standard curve. Virus titer was determined *via* plaque assay.

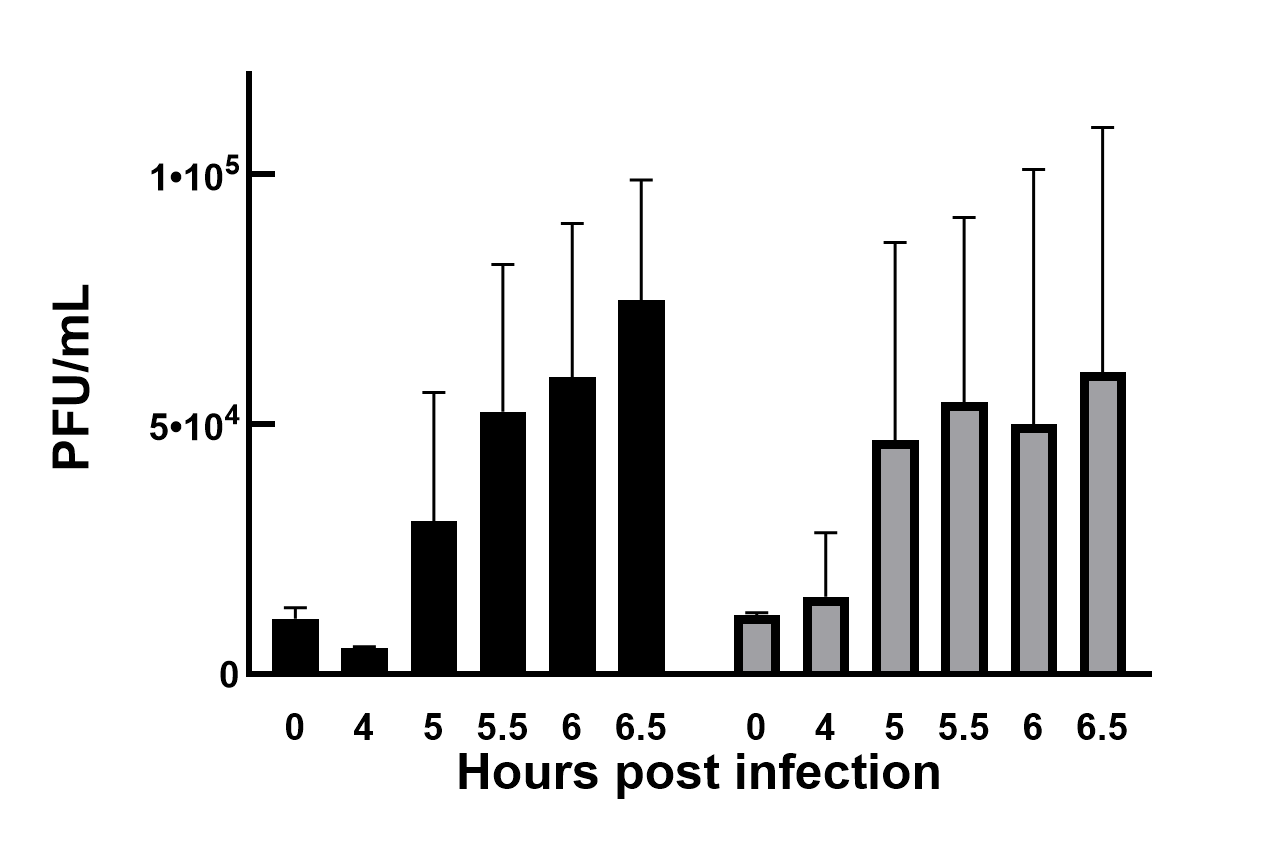

**Supplementary Figure 4.** Adsorption of ΨM2 virus particles to its host *M. marburgensis* between 4 h and 6.5 h post-infection in SAB medium (black) and MS medium (grey). The error bars represent the standard deviation of the assay, which was performed in biological replicates (n = 3).

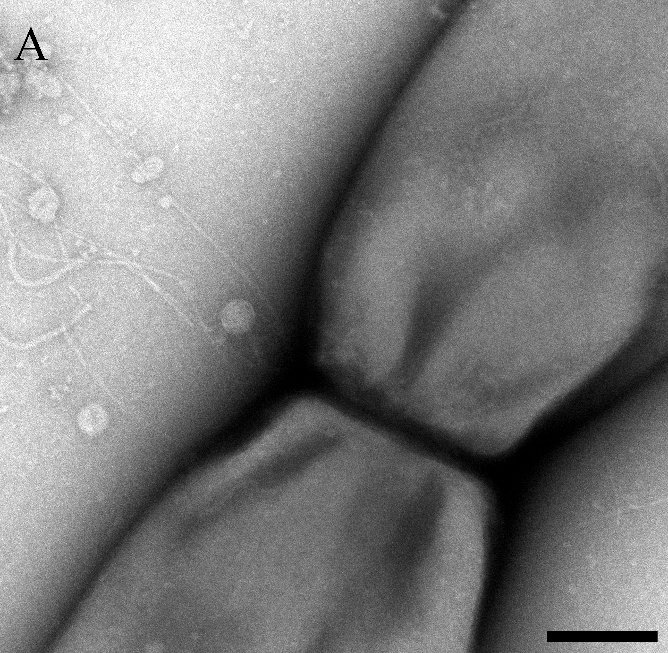

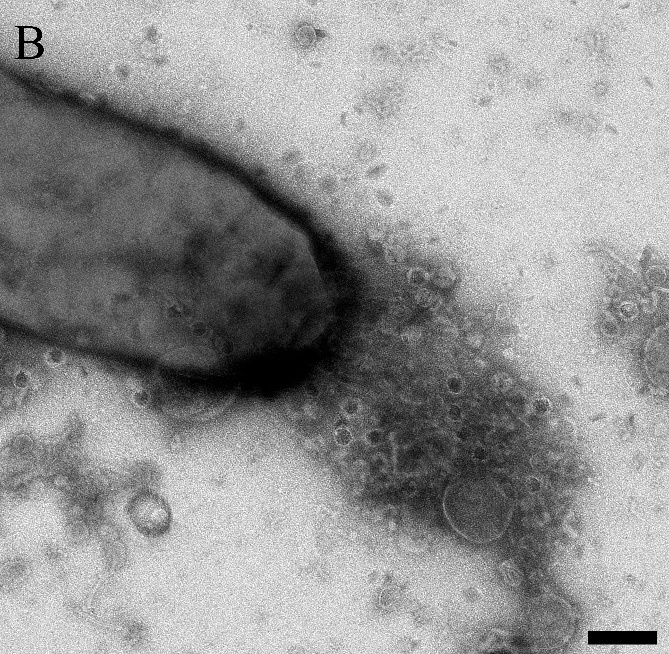

**Supplementary Figure 5.** TEM micrographs showing viral particles interacting with host cells near the division plane of daughter cells (A) and at a cell pole (B). The scale bars represent 200 nm.

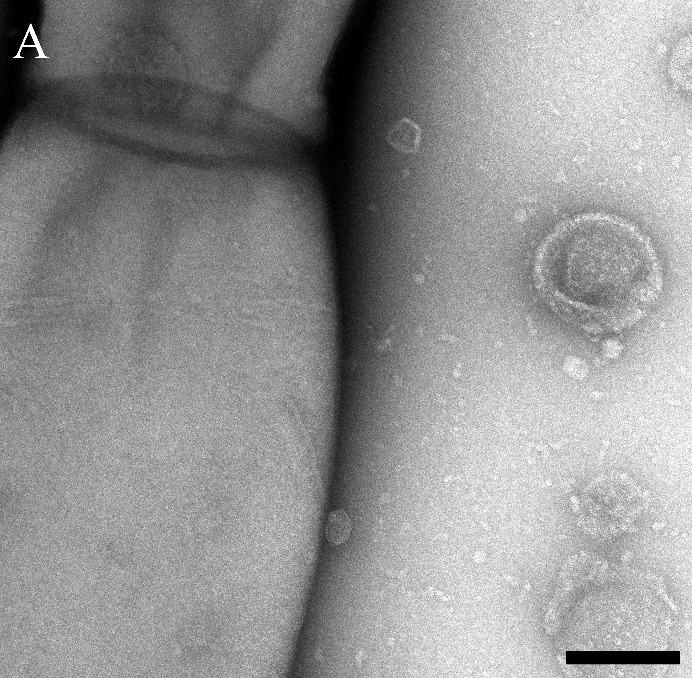

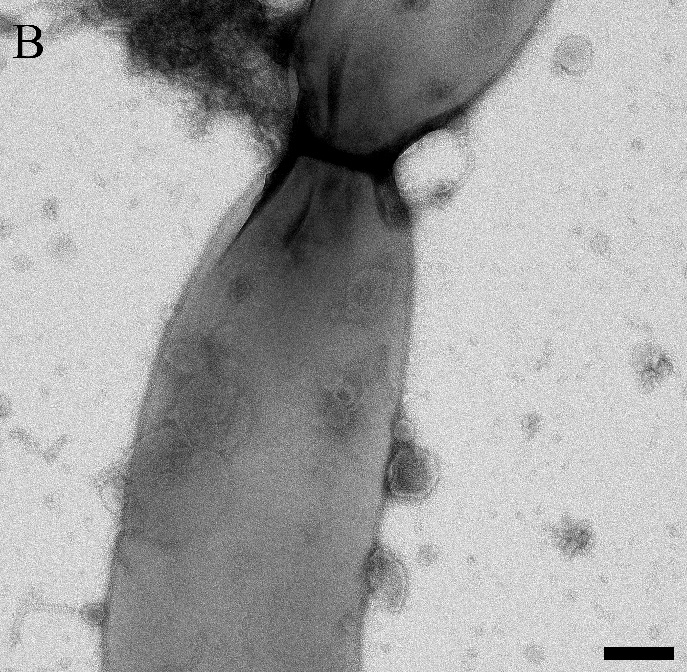

**Supplementary Figure 6.** TEM micrographs show viral particles attached to the host cell with their heads and tails near the cell wall (A) and with their heads but not their tails (B). The scale bars represent 200 nm.

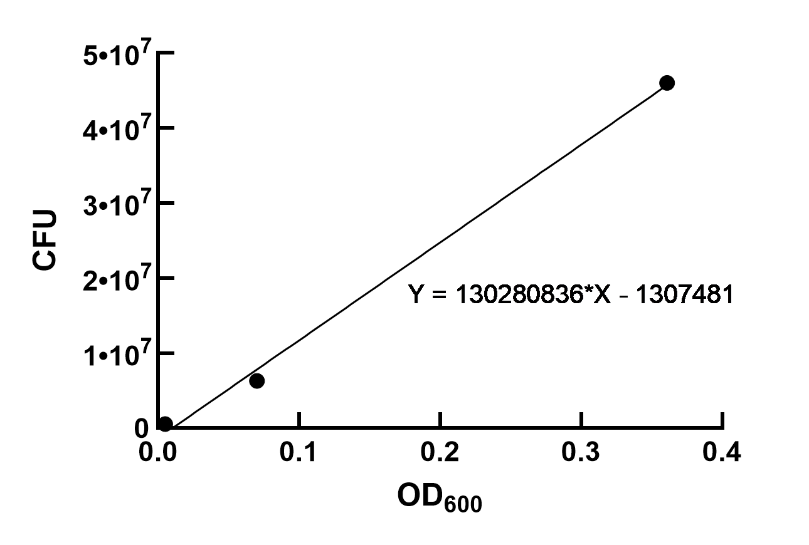

**Supplementary Figure 7.** Graph of plating experiment showing *M. marburgensis* CFU at different OD_600_ culture densities. A linear regression was performed and the resulting equation used for CFU calculation during experiments.

**Supplementary Table 1.** Sequencing data information provided as raw data and after trimming and filtering. The samples represent gDNA isolated from ΨM2-infected M. marburgensis cultures in MS medium taken at timepoints early exponential growth (A) and early stationary growth (B), as well as ΨM2-infected M. marburgensis cultures in SAB medium taken at timepoints early exponential growth (D), early stationary growth (E), and late stationary growth (F). Sample G represents a control culture of uninfected M. marburgensis in MS medium grown to early exponential growth.

| **Raw data** | | | | | |
| --- | --- | --- | --- | --- | --- |
| **Sample** | **Barcode** | **No. reads** | **No. bases** | **Mean length (bp)** | **Mean Q-score (Phred)** |
| **A** | barcode01 | 390762 | 612887945 | 1568.4 | 15.3 |
| **B** | barcode02 | 1955333 | 2909341173 | 1487.9 | 16 |
| **D** | barcode04 | 1383622 | 2860520101 | 2067.4 | 16.3 |
| **E** | barcode05 | 2255162 | 5575616308 | 2472.4 | 16.7 |
| **F** | barcode06 | 1176830 | 2502352502 | 2126.4 | 16.3 |
| **G** | barcode07 | 1395950 | 2504626001 | 1794.2 | 16.1 |
| **Nuclease-free water** | barcode09 | 266070 | 23057199 | 86.7 | 13.7 |
| **Trimmed and filtered data** | | | | | |
| **Sample** | **Barcode** | **No. reads** | **No. bases** | **Mean length (bp)** | **Mean Q-score (Phred)** |
| **A** | 135374 | 441131981 | 3258.6 | 20.4 | 135374 |
| **B** | 78992 | 500001509 | 6329.8 | 21.1 | 78992 |
| **D** | 67128 | 500004514 | 7448.5 | 21.7 | 67128 |
| **E** | 50014 | 500007608 | 9997.4 | 21.6 | 50014 |
| **F** | 70712 | 500005325 | 7071 | 21.5 | 70712 |
| **G** | 75251 | 500004259 | 6644.5 | 21.5 | 75251 |
| **Nuclease-free water** | 55 | 1175151 | 21366.4 | 19.3 | 55 |

**Supplementary Table 2.** Assembly metrics from Quast for the samples A, B, D, E, F, G.

| **Assembly metrics** | | | | | | | | |
| --- | --- | --- | --- | --- | --- | --- | --- | --- |
| **Sample** | **Genome fraction (%)** | **No. contigs** | **N50 (Mb)** | **Reference length (bp)** | **Total length (bp)** | **Largest contig (bp)** | **Reference GC (%)** | **Assembly GC (%)** |
| A | 100 | 3 | 1634695 | 1639146 | 1674987 | 1634695 | 26972 | 48.58 |
| B | 100 | 2 | 1634695 | 1639146 | 1639135 | 1634695 | 0 | 48.64 |
| D | 100 | 3 | 1634695 | 1639146 | 1674987 | 1634695 | 26972 | 48.58 |
| E | 100 | 3 | 1634695 | 1639146 | 1674987 | 1634695 | 26972 | 48.58 |
| F | 100 | 3 | 1634693 | 1639146 | 1670545 | 1634693 | 26972 | 48.58 |
| G | 100 | 2 | 1634693 | 1639146 | 1639762 | 1634693 | 0 | 48.64 |

**Supplementary Table 3.** BLAST results for plasmid pME2001. Results for contigs from the samples A, B, D, E, F, and G, which aligned with subject ID pME2001_plasmid_X17205.1_Methanobacterium_thermautotrophicum.

| **BLAST results pME2001** | | | | | | | | | | | |
| --- | --- | --- | --- | --- | --- | --- | --- | --- | --- | --- | --- |
| **Sample** | **Query ID** | **Identity** | **Aligned length (bp)** | **Mismatch** | **Gap open** | **Query start** | **Query end** | **Subject start** | **Subject end** | **E-value** | **Bit score** |
| A | contig_2 | 99.977 | 4440 | 0 | 1 | 2221 | 6660 | 1 | 4439 | 0 | 8192 |
| A | contig_2 | 99.977 | 4440 | 0 | 1 | 6661 | 11100 | 1 | 4439 | 0 | 8192 |
| A | contig_2 | 100 | 2220 | 0 | 0 | 1 | 2220 | 2220 | 4439 | 0 | 4100 |
| A | contig_2 | 99.955 | 2220 | 0 | 1 | 11101 | 13320 | 1 | 2219 | 0 | 4093 |
| B | contig_2 | 99.975 | 4006 | 0 | 1 | 1 | 4006 | 435 | 4439 | 0 | 7391 |
| B | contig_2 | 100 | 434 | 0 | 0 | 4007 | 4440 | 1 | 434 | 0 | 802 |
| D | contig_3 | 99.977 | 4440 | 0 | 1 | 6665 | 11104 | 4439 | 1 | 0 | 8192 |
| D | contig_3 | 99.977 | 4440 | 0 | 1 | 2225 | 6664 | 4439 | 1 | 0 | 8192 |
| D | contig_3 | 99.955 | 2224 | 0 | 1 | 1 | 2224 | 2223 | 1 | 0 | 4100 |
| D | contig_3 | 100 | 2216 | 0 | 0 | 11105 | 13320 | 4439 | 2224 | 0 | 4093 |
| E | contig_1 | 99.977 | 4440 | 0 | 1 | 3816 | 8255 | 1 | 4439 | 0 | 8192 |
| E | contig_1 | 99.977 | 4440 | 0 | 1 | 8256 | 12695 | 1 | 4439 | 0 | 8192 |
| E | contig_1 | 99.974 | 3815 | 0 | 1 | 1 | 3815 | 626 | 4439 | 0 | 7038 |
| E | contig_1 | 100 | 625 | 0 | 0 | 12696 | 13320 | 1 | 625 | 0 | 1155 |
| F | contig_3 | 99.977 | 4440 | 0 | 1 | 667 | 5106 | 1 | 4439 | 0 | 8192 |
| F | contig_3 | 99.974 | 3774 | 0 | 1 | 5107 | 8880 | 1 | 3773 | 0 | 6962 |
| F | contig_3 | 100 | 666 | 0 | 0 | 1 | 666 | 3774 | 4439 | 0 | 1230 |
| G | contig_1 | 100 | 629 | 0 | 0 | 1 | 629 | 2011 | 2639 | 0 | 1162 |
| G | contig_3 | 99.956 | 2258 | 0 | 1 | 2183 | 4440 | 1 | 2257 | 0 | 4163 |
| G | contig_3 | 100 | 2182 | 0 | 0 | 1 | 2182 | 2258 | 4439 | 0 | 4030 |

**Supplementary Table 4.** GeNomad output for assembled genome data from samples A, D, E, and F, which contained contigs that were recognized as viral by the program.

| **GeNomad output** | | | | |
| --- | --- | --- | --- | --- |
| **Sample** | **A** | **D** | **E** | **F** |
| **Seq_name** | contig_3 | contig_2 | contig_3 | contig_2 |
| **Length (bp)** | 26,972 | 26,972 | 26,972 | 26,972 |
| **Topology** | No terminal repeats | No terminal repeats | No terminal repeats | No terminal repeats |
| **Coordinates** | NA | NA | NA | NA |
| **n_genes** | 41 | 41 | 41 | 40 |
| **Genetic_code** | 11 | 11 | 11 | 11 |
| **Virus_score** | 0.995 | 0.9972 | 0.9945 | 0.9975 |
| **fdr** | 0.005 | 0.0028 | 0.0055 | 0.0025 |
| **n_hallmarks** | 6 | 5 | 5 | 5 |
| **Marker_enrichment** | 33.2194 | 31.7314 | 33.4497 | 31.7314 |
| **Taxonomy** | Viruses; Duplodnaviria; Heunggongvirae; Uroviricota; Caudoviricetes; Methanobavirales; Leisingerviridae | Viruses; Duplodnaviria; Heunggongvirae; Uroviricota; Caudoviricetes; Methanobavirales; Leisingerviridae | Viruses; Duplodnaviria; Heunggongvirae; Uroviricota; Caudoviricetes; Methanobavirales; Leisingerviridae | Viruses; Duplodnaviria; Heunggongvirae; Uroviricota; Caudoviricetes; Methanobavirales; Leisingerviridae |

**Supplementary Table 5.** VIBRANT output for assembled genome data from samples A, D, E, and F, which contained contigs that were recognized as viral by the program.

| **VIBRANT output** | | | | |
| --- | --- | --- | --- | --- |
| **Sample** | **A** | **D** | **E** | **F** |
| **Scaffold** | contig_3 | contig_2 | contig_3 | contig_2 |
| **Total genes** | 41 | 41 | 41 | 40 |
| **All KEGG** | 2 | 2 | 2 | 2 |
| **KEGG v-score** | 7.37 | 7.37 | 7.37 | 7.37 |
| **All Pfam** | 6 | 5 | 5 | 5 |
| **Pfam v-score** | 19.14 | 16.29 | 16.29 | 16.29 |
| **All VOG** | 32 | 32 | 32 | 31 |
| **VOG v-score** | 39.9 | 29.92 | 29.92 | 29.9 |
| **KEGG int-rep** | 1 | 1 | 1 | 1 |
| **KEGG zero** | 0 | 0 | 0 | 0 |
| **Pfam int-rep** | 1 | 1 | 1 | 1 |
| **Pfam zero** | 0 | 0 | 0 | 0 |
| **VOG redoxin** | 0 | 0 | 0 | 0 |
| **VOG rec-tran** | 0 | 0 | 0 | 0 |
| **VOG int** | 0 | 0 | 0 | 0 |
| **VOG RnR** | 0 | 0 | 0 | 0 |
| **VOG DNA** | 0 | 0 | 0 | 0 |
| **KEGG restriction check** | 0 | 0 | 0 | 0 |
| **KEGG toxin check** | 0 | 0 | 0 | 0 |
| **VOG special** | 7 | 6 | 6 | 6 |
| **Annotation check** | 2 | 2 | 2 | 2 |
| **p_v check** | 4 | 3 | 3 | 3 |
| **p_k check** | 0 | 0 | 0 | 0 |
| **k_v check** | 0 | 0 | 0 | 0 |
| **k check** | 0 | 0 | 0 | 0 |
| **p check** | 0 | 0 | 0 | 0 |
| **v check** | 26 | 27 | 27 | 26 |
| **h check** | 9 | 9 | 9 | 9 |

**Supplementary Table 6.** BLAST results for ΨM2 virus genome. Results for contigs from the samples A, D, E, and F, which aligned with subject ID psiM2_phage_Methanobacterium_NC_001902.1.

| **BLAST results ΨM2 virus genome** | | | | | | | | | | | |
| --- | --- | --- | --- | --- | --- | --- | --- | --- | --- | --- | --- |
| **Sample** | **Query ID** | **Identity** | **Aligned length (bp)** | **Mismatch** | **Gap open** | **Query start** | **Query end** | **Subject start** | **Subject end** | **E-value** | **Bit score** |
| A | contig_3 | 99.973 | 11023 | 2 | 1 | 15951 | 26972 | 16760 | 5738 | 0 | 20338 |
| A | contig_3 | 99.983 | 5960 | 1 | 0 | 10112 | 16071 | 22554 | 16595 | 0 | 11001 |
| A | contig_3 | 98.273 | 4110 | 69 | 2 | 5863 | 9971 | 26111 | 22003 | 0 | 7195 |
| A | contig_3 | 99.968 | 3116 | 0 | 1 | 1 | 3116 | 5737 | 2623 | 0 | 5747 |
| A | contig_3 | 99.929 | 2814 | 1 | 1 | 3050 | 5862 | 2814 | 1 | 0 | 5184 |
| A | contig_3 | 100 | 67 | 0 | 0 | 2925 | 2991 | 2689 | 2623 | 1.84E-29 | 124 |
| A | contig_3 | 100 | 67 | 0 | 0 | 3175 | 3241 | 2814 | 2748 | 1.84E-29 | 124 |
| A | contig_3 | 100 | 54 | 0 | 0 | 8929 | 8982 | 22799 | 22746 | 3.10E-22 | 100 |
| A | contig_3 | 100 | 54 | 0 | 0 | 9175 | 9228 | 23045 | 22992 | 3.10E-22 | 100 |
| D | contig_2 | 99.972 | 14139 | 2 | 2 | 9100 | 23237 | 2623 | 16760 | 0 | 26086 |
| D | contig_2 | 98.273 | 4110 | 69 | 2 | 2245 | 6353 | 22003 | 26111 | 0 | 7195 |
| D | contig_2 | 99.974 | 3856 | 1 | 0 | 23117 | 26972 | 16595 | 20450 | 0 | 7116 |
| D | contig_2 | 99.929 | 2814 | 1 | 1 | 6354 | 9166 | 1 | 2814 | 0 | 5184 |
| D | contig_2 | 100 | 2104 | 0 | 0 | 1 | 2104 | 20451 | 22554 | 0 | 3886 |
| D | contig_2 | 100 | 67 | 0 | 0 | 9225 | 9291 | 2623 | 2689 | 1.84E-29 | 124 |
| D | contig_2 | 100 | 67 | 0 | 0 | 8975 | 9041 | 2748 | 2814 | 1.84E-29 | 124 |
| D | contig_2 | 100 | 54 | 0 | 0 | 3234 | 3287 | 22746 | 22799 | 3.10E-22 | 100 |
| D | contig_2 | 100 | 54 | 0 | 0 | 2988 | 3041 | 22992 | 23045 | 3.10E-22 | 100 |
| E | contig_3 | 99.972 | 14139 | 2 | 2 | 11575 | 25712 | 16760 | 2623 | 0 | 26086 |
| E | contig_3 | 99.983 | 5960 | 1 | 0 | 5736 | 11695 | 22554 | 16595 | 0 | 11001 |
| E | contig_3 | 98.273 | 4110 | 69 | 2 | 1487 | 5595 | 26111 | 22003 | 0 | 7195 |
| E | contig_3 | 99.866 | 1487 | 1 | 1 | 1 | 1486 | 1487 | 1 | 0 | 2734 |
| E | contig_3 | 100 | 1327 | 0 | 0 | 25646 | 26972 | 2814 | 1488 | 0 | 2451 |
| E | contig_3 | 100 | 67 | 0 | 0 | 25521 | 25587 | 2689 | 2623 | 1.84E-29 | 124 |
| E | contig_3 | 100 | 67 | 0 | 0 | 25771 | 25837 | 2814 | 2748 | 1.84E-29 | 124 |
| E | contig_3 | 100 | 54 | 0 | 0 | 4553 | 4606 | 22799 | 22746 | 3.10E-22 | 100 |
| E | contig_3 | 100 | 54 | 0 | 0 | 4799 | 4852 | 23045 | 22992 | 3.10E-22 | 100 |
| F | contig_2 | 99.979 | 14139 | 1 | 2 | 7003 | 21140 | 2623 | 16760 | 0 | 26092 |
| F | contig_2 | 99.983 | 5953 | 1 | 0 | 21020 | 26972 | 16595 | 22547 | 0 | 10988 |
| F | contig_2 | 98.273 | 4110 | 69 | 2 | 148 | 4256 | 22003 | 26111 | 0 | 7195 |
| F | contig_2 | 99.929 | 2814 | 1 | 1 | 4257 | 7069 | 1 | 2814 | 0 | 5184 |
| F | contig_2 | 100 | 67 | 0 | 0 | 7128 | 7194 | 2623 | 2689 | 1.84E-29 | 124 |
| F | contig_2 | 100 | 67 | 0 | 0 | 6878 | 6944 | 2748 | 2814 | 1.84E-29 | 124 |
| F | contig_2 | 100 | 54 | 0 | 0 | 1137 | 1190 | 22746 | 22799 | 3.10E-22 | 100 |
| F | contig_2 | 100 | 54 | 0 | 0 | 891 | 944 | 22992 | 23045 | 3.10E-22 | 100 |

**Supplementary Table 7.** Insertions identified through manual curation in Geneious Prime, including their genomic position, size, predicted annotation of the open reading frame (ORF) in which they occur, and their frequency within sequencing reads (%).

| **Position (bp)** | **Size (bp)** | **ORF/intergenic** | **Protein annotation** | **Frequency within reads (%)** | | | | |
| --- | --- | --- | --- | --- | --- | --- | --- | --- |
|  |  |  |  | **Sample A** | **Sample B** | **Sample D** | **Sample E** | **Sample F** |
| 2622 | >124 | integenic | - | 0.36 | 0.26 | 0.33 | 0.35 | 0.34 |
| 3260 | 1 | ORF | ribonuclease | 0.93 | 0.92 | 0.93 | 0.94 | 0.93 |
| 16608 | >15 | ORF | tape measure | 0.63 | 0.57 | 0.65 | 0.65 | 0.62 |
| 22005 | >30 | ORF | spike protein | 0.16 | 0.15 | 0.13 | 0.13 | 0.14 |
| 22229 | 1 | ORF | spike protein | 0.20 | 0.20 | 0.17 | 0.17 | 0.18 |
| 22246 | 1 | ORF | spike protein | 0.16 | 0.15 | 0.13 | 0.12 | 0.13 |
| 22472 | 692 | ORF | spike protein / PSIM1 variant | 0.20 | 0.15 | 0.19 | 0.21 | 0.20 |

**Supplementary Table 8.** Single-nucleotide variants (SNVs) identified using inStrain, including the reference and variant nucleotides, codon position, resulting amino acid change, and their frequency within sequencing reads (%).

| **Position (bp)** | **Nucleotide** | | **Codon position** | **Amino Acid** | | **Frequency within reads (%)** | | | | |
| --- | --- | --- | --- | --- | --- | --- | --- | --- | --- | --- |
|  | **Reference** | **Variant** |  | **Reference** | **Variant** | **Sample A** | **Sample B** | **Sample D** | **Sample E** | **Sample F** |
| 10291 | C | T | 3 | V | V | 0.44 | 0.48 | 0.43 | 0.44 | 0.43 |
| 22137 | C | T | 1 | P | S | 0.13 | 0.11 | 0.11 | 0.12 | 0.13 |
| 22140 | A | C | 1 | T | P | 0.12 | 0.11 | 0.11 | 0.12 | 0.13 |
| 22144 | C | A | 2 | S | Y | 0.12 | 0.10 | 0.10 | 0.11 | 0.12 |
| 22145 | T | C | 3 | S | S | 0.12 | 0.11 | 0.11 | 0.11 | 0.12 |
| 22147 | G | C | 2 | W | S | 0.13 | 0.11 | 0.11 | 0.12 | 0.13 |
| 22148 | G | A | 3 | W | * | 0.16 | 0.14 | 0.13 | 0.15 | 0.16 |
| 22163 | C | A | 3 | I | I | 0.15 | 0.14 | 0.14 | 0.15 | 0.16 |
| 22164 | T | A | 1 | L | M | 0.15 | 0.14 | 0.14 | 0.15 | 0.16 |
| 22168 | A | G | 2 | Y | C | 0.15 | 0.14 | 0.14 | 0.15 | 0.16 |
| 22169 | T | C | 3 | Y | Y | 0.15 | 0.14 | 0.14 | 0.15 | 0.16 |
| 22175 | T | C | 3 | N | N | 0.15 | 0.14 | 0.14 | 0.15 | 0.16 |
| 22183 | C | G | 2 | T | S | 0.13 | 0.13 | 0.12 | 0.13 | 0.14 |
| 22184 | T | A | 3 | T | T | 0.14 | 0.13 | 0.12 | 0.13 | 0.14 |
| 22187 | A | G | 3 | G | G | 0.14 | 0.14 | 0.14 | 0.15 | 0.16 |
| 22188 | T | G | 1 | S | A | 0.13 | 0.14 | 0.14 | 0.15 | 0.16 |
| 22198 | T | C | 2 | I | T | 0.13 | 0.12 | 0.12 | 0.13 | 0.13 |
| 22199 | C | G | 3 | I | M | 0.14 | 0.12 | 0.11 | 0.12 | 0.13 |
| 22202 | C | T | 3 | C | C | 0.12 | 0.12 | 0.11 | 0.12 | 0.12 |
| 22212 | C | G | 1 | H | D | 0.14 | 0.13 | 0.12 | 0.13 | 0.14 |
| 22213 | A | C | 2 | H | P | 0.15 | 0.14 | 0.13 | 0.14 | 0.15 |
| 22218 | G | A | 1 | D | N | 0.17 | 0.15 | 0.14 | 0.16 | 0.17 |
| 22225 | T | G | 2 | V | G | 0.16 | 0.15 | 0.14 | 0.16 | 0.17 |
| 22229 | C | T | 3 | G | G | 0.15 | 0.14 | 0.14 | 0.15 | 0.16 |
| 22230 | A | T | 1 | T | S | 0.16 | 0.14 | 0.14 | 0.15 | 0.16 |
| 22231 | C | T | 2 | T | I | 0.16 | 0.14 | 0.13 | 0.15 | 0.15 |
| 22234 | C | T | 2 | S | F | 0.15 | 0.14 | 0.14 | 0.15 | 0.15 |
| 22244 | C | G | 3 | T | T | 0.12 | 0.11 | 0.11 | 0.11 | 0.12 |
| 22253 | C | A | 3 | P | P | 0.19 | 0.17 | 0.17 | 0.19 | 0.19 |
| 22266 | A | C | 1 | N | H | 0.14 | 0.13 | 0.14 | 0.15 | 0.15 |
| 22269 | T | C | 1 | L | L | 0.18 | 0.16 | 0.15 | 0.17 | 0.18 |
| 22271 | G | T | 3 | L | F | 0.18 | 0.15 | 0.15 | 0.16 | 0.17 |
| 22274 | T | C | 3 | C | C | 0.17 | 0.15 | 0.14 | 0.15 | 0.16 |
| 22287 | A | T | 1 | R | * | 0.22 | 0.16 | 0.16 | 0.17 | 0.18 |
| 22295 | C | T | 3 | R | R | 0.23 | 0.19 | 0.19 | 0.20 | 0.21 |
| 22296 | G | A | 1 | A | T | 0.23 | 0.20 | 0.20 | 0.21 | 0.22 |
| 22301 | A | G | 3 | L | L | 0.22 | 0.21 | 0.20 | 0.22 | 0.23 |
| 22307 | A | G | 3 | E | E | 0.23 | 0.19 | 0.20 | 0.20 | 0.22 |
| 22310 | C | A | 3 | G | G | 0.21 | 0.19 | 0.19 | 0.20 | 0.21 |
| 22313 | T | C | 3 | Y | Y | 0.21 | 0.19 | 0.19 | 0.20 | 0.22 |
| 22316 | A | C | 3 | A | A | 0.21 | 0.20 | 0.20 | 0.21 | 0.22 |
| 22332 | C | G | 1 | P | A | 0.14 | 0.12 | 0.13 | 0.13 | 0.14 |
| 22333 | C | T | 2 | P | L | 0.14 | 0.12 | 0.13 | 0.13 | 0.14 |
| 22334 | T | G | 3 | P | P | 0.13 | 0.11 | 0.11 | 0.12 | 0.13 |
| 22340 | T | G | 3 | G | G | 0.24 | 0.21 | 0.22 | 0.22 | 0.23 |
| 22343 | T | C | 3 | Y | Y | 0.26 | 0.20 | 0.22 | 0.22 | 0.23 |
| 22350 | G | A | 1 | A | T | 0.25 | 0.21 | 0.22 | 0.22 | 0.23 |
| 22435 | A | C | 2 | E | A | 0.38 | 0.31 | 0.30 | 0.31 | 0.33 |
| 22467 | G | A | 1 | G | R | 0.37 | 0.29 | 0.31 | 0.29 | 0.32 |
| 22471 | T | A | 2 | L | Q | 0.33 | 0.24 | 0.27 | 0.25 | 0.27 |

**Supplementary Table 9.** Overview of the concentration of free virus particles (PFU/mL) measured via plaque assay from samples of the adsorption assay taken at the time of infection (0 h) and 1 h post-infection. The adsorption rate was calculated with the formula k = 2.3 / B * t × log10(P(0)/P(t)) with host cell concentration (B), free virus concentration (P) at time point 0 and time point of adsorption (t).

| Medium | Replicate | 0h | 1h | Host cell  concentration | Adsorption rate | Mean adsorption rate |
| --- | --- | --- | --- | --- | --- | --- |
| SAB | 1 | 25000 | 18400 | 600000 | 4.8 * 10^-7^ | 3.5 * 10^-7^ |
|  | 2 | 31400 | 17600 | 600000 | 2.5 * 10^-7^ |  |
|  | 3 | 28000 | 17400 | 600000 | 3.1 * 10^-7^ |  |
| MS | 1 | 25200 | 19000 | 800000 | 3.9 * 10^-7^ | 3.8 * 10^-7^ |
|  | 2 | 39600 | 19000 | 1000000 | 1.2 * 10^-7^ |  |
|  | 3 | 21000 | 17600 | 800000 | 6.2 * 10^-7^ |  |

**Supplementary Table 10.** Overview of virus particle sizes measured from 10 imaged ΨM2 virus particles.

| Measurement (nm) | Head width | Head height | Tail width | Tail length | Number of tail segments |
| --- | --- | --- | --- | --- | --- |
| Image A | 61.303 | 61.432 | 14.834 | 244.829 | 52 |
| Image B | 57.407 | 62.465 | 13.485 | 250.495 | 50 |
| Image C | 68.432 | 76.720 | 12.414 | 233.068 | 51 |
| Image D | 55.384 | 60.668 | 12.149 | 237.612 | 50 |
| Image E | 44.508 | 48.016 | 11.563 | 223.891 | 48 |
| Image F | 55.775 | 57.497 | 11.990 | 231.988 | 52 |
| Image G | 63.089 | 64.625 | 11.277 | 235.506 | - |
| Image H | 64.472 | 64.105 | 12.026 | 225.306 | - |
| Image I | 65.045 | 62.118 | 11.763 | 237.218 | 49 |
| Image J | 59.140 | 62.361 | 11.535 | 236.874 | 50 |
| Mean | 59.456 | 62.001 | 12.304 | 235.679 | 50.25 |
| Standard deviation | 6.414 | 6.680 | 1.0245 | 7.601 | 1.299 |

**Supplementary Result 1.**

**Different media supplements affect plaque size and appearance**

To further investigate the effects of medium composition on virus-host interaction, we followed up on an observation from a previous study, which mentioned differing plaque formation depending on media complexity (Meile et al., 1989). The authors had performed plaque assays and observed smaller plaques on complete medium compared to plaques on minimal medium. They used a mineral salt medium as the minimal medium and added 2 g/L each of yeast extract, acetate, and formate to make complete medium (Schönheit et al., 1980). For plates, they also added 1 mM titanium(III) citrate (Meile et al., 1989). We repeated the plaque assay on solid MS medium supplemented only with yeast extract. The plaques on standard MS medium plates displayed larger diameters and sharp edges, while plaques on the MS medium plates supplemented with yeast extract were smaller in diameter and had plaque halos (areas of thinner host cell growth) around them (**Supplementary Figure 1**). The median plaque diameter with (0.67 mm, n = 49) and without (1.41 mm, n = 36) yeast extract supplementation differed significantly (Mann-Whitney U = 112, p < 0.0001 two-tailed). In addition, when MS medium plates without yeast extract were imaged after less than 24 hours of incubation, large plaques with *M. marburgensis* colonies in the middle could be observed (**Supplementary Figure 2**).
